# Dynamic calcium-mediated stress response and recovery signatures in the fungal pathogen, *Candida albicans*

**DOI:** 10.1101/2023.04.20.537637

**Authors:** CV Giuraniuc, C Parkin, MC Almeida, M Fricker, P Shadmani, S Nye, S Wehmeier, S Chawla, T Bedekovic, L Lehtovirta-Morley, D Richards, NA Gow, AC Brand

**Affiliations:** MRC Centre for Medical Mycology at the University of Exeter, Exeter, UK; Living Systems Institute, University of Exeter, UK; School of Plant Sciences, University of Oxford, Oxford UK; School of Medicine, Medical Sciences & Nutrition, University of Aberdeen, UK; Department of Physics and Astronomy, University of Exeter, UK

**Author notes:** Joint first authors (CV Giuraniuc, C Parkin).

## Abstract

Calcium (Ca^2+^) is an important second messenger for activating stress response signalling and cell adaptation in eukaryotic cells yet intracellular Ca^2+^-dynamics in fungi is poorly understood due to lack of effective real-time Ca^2+^ reporters. We engineered the GCaMP6f construct for use in the fungal pathogen, *Candida albicans*, and used live-cell imaging to observe dynamic Ca^2+^ spiking as well as slower changes in ambient Ca^2+^-GCaMP levels elicited by stress or gene deletion. Short-term exposure to membrane, osmotic or oxidative stress generated immediate stress-specific responses and repeated exposure revealed differential recovery signatures. Osmotic stress caused yeast cell shrinkage and no adaptation response, where Ca^2+^-GCaMP spiking was inhibited by 1 M NaCl but not by 0.66 M CaCl_2._ Treatment with SDS caused a spike-burst, raised ambient Ca^2+^-GCaMP levels and significant cell death, but surviving cells adapted over subsequent exposures. Treatment with 5 mM H_2_O_2_ abolished spiking and caused transient autofluorescence but cells adapted such that spiking returned and autofluorescence diminished on repeated exposure. Adaptation to H_2_O_2_ was dependent on Cap1, extracellular Ca^2+^ and calcineurin, but not on its downstream target, Crz1. Ca^2+^-dynamics were not affected by H_2_O_2_ in the *hog1*Δ or *yvc1*Δ mutants, suggesting a pre-adapted, resistant state, possibly due to changes in membrane permeability. Live-cell imaging of Ca^2+^-GCaMP responses in individual cells has therefore revealed the dynamics of Ca^2+^-influx, signalling and homeostasis and their role in the temporal stress response signatures of *C. albicans*.

## Introduction

*Candida albicans* is an opportunistic fungal pathogen that causes around 400,000 life-threatening bloodstream infections a year in patients undergoing immunosuppressive treatments, in addition to mucosal infections in millions of predisposed patients, particularly women of child-bearing age (Brown et al., 2012, Kullberg and Arendrup, 2015). The regulation of mechanisms that contribute to pathogenesis by countering stress within the host environment or by the action of antifungal drugs have been the focus of numerous studies. Calcium (Ca^2+^) is an essential trace element in all eukaryotic cells, where it acts as an enzyme co-factor but is also an important second messenger that activates cell stress responses. In *C. albicans*, this includes Ca^2+^ and membrane stress, which are key to its survival in serum and resistance to membrane-targeting antifungal drugs (Blankenship and Heitman, 2005, Cruz et al., 2002, Sanglard et al., 2003). Cytosolic [Ca^2+^]_cyt_ is maintained at ∼ 80-120 nM (Iida et al., 1990) such that even small changes due to influx across the plasma-membrane or release from intracellular stores acts as an immediate signal. Ca^2+^– binding to calmodulin activates the calcineurin phosphatase that has many effectors but downstream activation of Crz1-dependent gene expression is the best-characterised to date (Karababa et al., 2006).

Despite its importance in resilience to stress, the dynamics and regulation of Ca^2+^ flux in *C. albicans*, and indeed any fungus, are poorly understood due to the lack of effective tools with which to investigate changes both in real time and at the level of single cells. Studies have therefore been limited to those at the population level. For example, the ^45^Ca^2+^ radioisotope was used to demonstrate that the plasma-membrane Cch1-Mid1 channel is involved in Ca^2+^ uptake, and the photoprotein, aequorin, was used to show that cells swiftly take up Ca^2+^ in response to the detergent, SDS (Brand et al., 2007, Sanglard, 2021). However, these methods yield no information on population heterogeneity, single-cell dynamics or longer-term Ca^2+^ response signatures.

We have developed a new genetically-encoded calcium indicator (GECI) based on the high signal-to-noise ratio, single-fluorophore indicator, GCaMP6 (Chen et al., 2013). The three variants of GCaMP6 (fast, medium and slow) were originally developed for use in neurons, which maintain cytoplasmic [Ca^2+^] at 65 – 70 nM and have fast Ca^2+^ dynamics (Sabatini et al., 2002, Chen et al., 2013). In *C. albicans*, cytoplasmic [Ca^2+^] is maintained at similar levels but the dynamics of cytoplasmic Ca^2+^ flux are completely unknown. We selected the GCaMP6f variant and used it to investigate Ca^2+^ responses in *C. albicans* wild-type and signalling-pathway mutant cells when subjected to repeated exposure to stress compounds. Ca^2+^-GCaMP6 activity in resting cells appeared as cytoplasmic spikes of 5 – 6 s in duration whose rate was dependent on extracellular [Ca^2+^] and an environmental pH of ≥ 7.0. In addition to spiking, some treatments raised the ambient level of cytoplasmic Ca^2+^-GCaMP, which was differentiated from changes in cell autofluorescence using an empty-vector control strain. Live-cell imaging of all three responses during treatment with cell stressors, including osmotic stress (1 M NaCl, 0.66 M CaCl_2_), membrane stress (0.05% SDS) and oxidative stress (5 mM H_2_O_2_), identified distinct stress-specific Ca^2+^-GCaMP6 signatures that suggest differential perturbation of plasma-membrane Ca^2+^-channel activity and intracellular Ca^2+^ homeostasis mechanisms. While osmotic stresses elicited the same response on each repeated treatment, Ca^2+^-dynamics slowly reverted to normal spiking and ambient levels in the presence of SDS or H_2_O_2_, suggesting that cells were able to adapt. By expressing GCaMP6 in mutant strains, we showed that the Cap1 transcription factor, extracelllar Ca^2+^ and the calcineurin phosphatase was required for adaptation to oxidative stress in *C. albicans* but the calcineurin-dependent transcription factor, Crz1 was not. Nevertheless, deletion of Crz1 appeared to disorganise intracellular Ca^2+^ homeostasis. In contrast to wild-type cells, Ca^2+^-GCaMP6 spiking was not affected in the mutant lacking Hog1, a MAPK in both osmotic and oxidative stress pathways, suggesting that this mutant was pre-adapted to this stress. The GCaMP6 reporter therefore provides a novel method with which to investigate the regulation of Ca^2+^ homeostasis, stress-signalling and adaptation over time, both at the population level and within individual cells.

## Results

### The GCaMP6 response in *C. albicans* is [Ca^2+^]_e_ and pH-dependent

The GCaMP6f (fast) variant was synthesised to incorporate circularly-permutated yEGFP3 along with the Ca^2+^-binding domain of calmodulin and the M13 peptide codon-optimised for *C. albicans* (Cormack et al., 1997, Chen et al., 2013) (Fig. S1). The construct was cloned into the CIpNAT and CIp10 plasmids (carrying nourseothricin resistance or the essential *URA3* gene as selectable markers, respectively) and transformed into gene deletion mutants and their relevant control strains (Table S2). GCaMP6 activity was imaged in yeast cells in a CellASIC microfluidics plate perfused with trace-metal-free Modified Soll’s Medium (Brand et al., 2007) supplemented with CaCl_2_ (see below). Cells were acclimatised by incubation at 30 °C without imaging. After 40 min, Stage 1 commenced with imaging in FITC and DIC every 5 s in baseline conditions (Fig. 1A, Movie S1). After 5 min, the perfusion channel was switched to MSM containing a compound of interest where cells were exposed for 10 min (Stage 2). This was followed by the Stage 3 washout step and reversion to baseline conditions for 15 min (Stage 3). Cells were imaged continuously from the start of Stage 1 to the end of Stage 3. In later experiments, the 3-stage process was repeated at microscope positions 2 and 3, which allowed quantification of the effect on a cell population of 3 exposures to the same treatment whilst minimising photo-stress (Fig 1B). Continued cell growth was confirmed by imaging in DIC only at a 4^th^ position without treatment. The imaging output of each cell per field of view over time was analysed by custom software (see Methods) and revealed two modes of Ca^2+^-GCaMP response: dynamic fluorescent cytoplasmic spiking (plotted as spikes/cell/min) and what we termed ‘ambient Ca^2+^-GCaMP level’, which was the Ca^2+^-GCaMP-dependent change in fluorescence output above Stage 1 values. The ambient Ca^2+^-GCaMP signal was differentiated from autofluorescence by comparing the fluorescence outputs between strains expressing either GCaMP or the empty-vector. Fast-acquisition imaging at 146 ms showed that spike duration was 5 – 6 s (Fig 1D) but imaging at this frame-rate caused photo-stress and significant cell death within 15 min. A frame rate of 5 s was therefore used as it permitted informative imaging at each microscope position for 30 min without affecting cell viability.

**Fig. 1.**
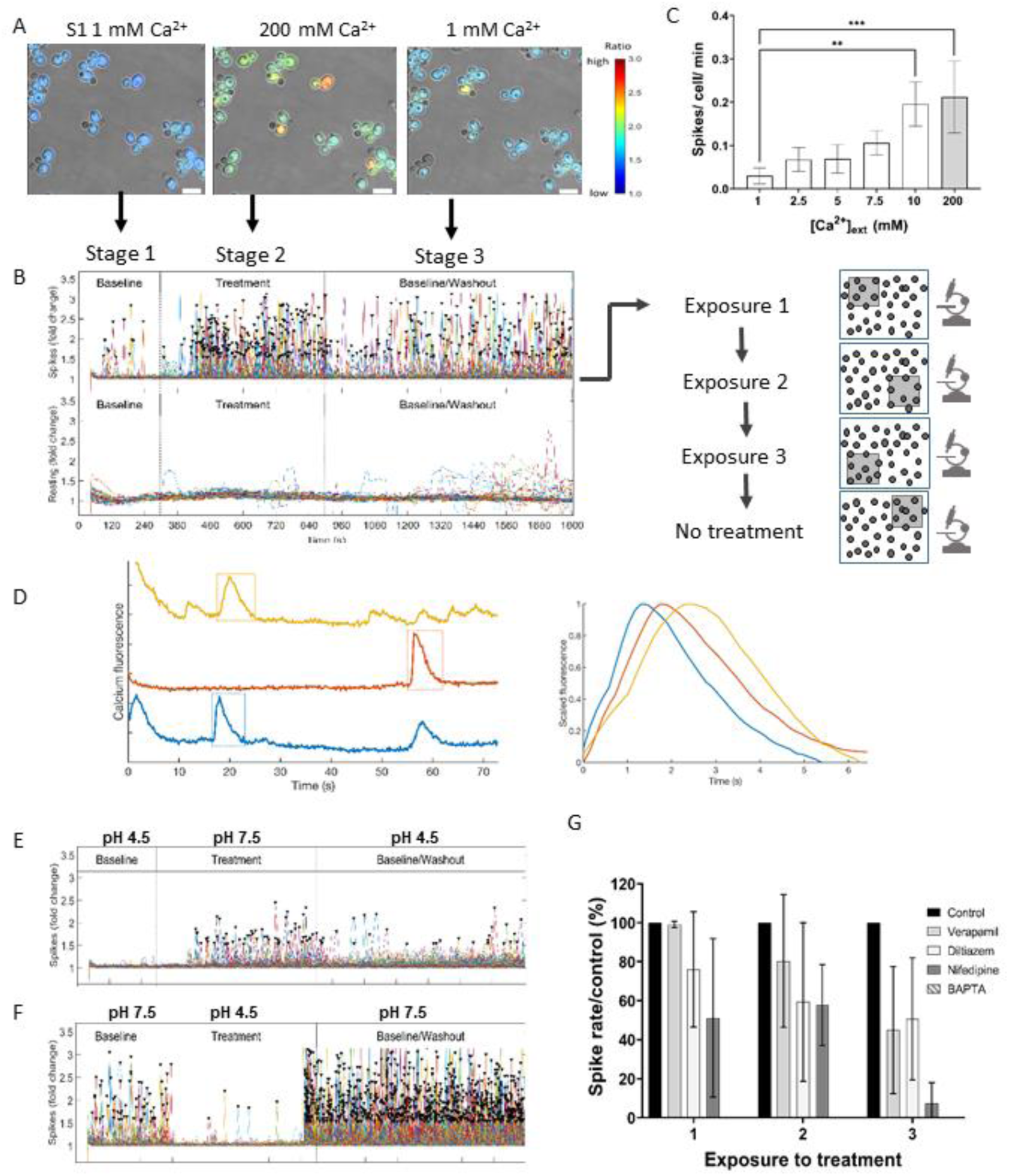
GCaMP6 spiking is dependent on [Ca^2+^]_ext_ and a neutral external pH. A. GCaMP imaging during Stage 1 (S1, 1 mM Ca^2+^), Stage 2 (S2, 200 mM Ca^2+^) and Stage 3 (S3,1 mM Ca^2+^). Bars = 10 um. B. Top panel – Ca^2+^-GCaMP spike distribution across the 3-stage time-course during Exposure 1. Bottom panel – baseline fluorescence signal over the same time-course. Cells were exposed to treatment 3 times, each at a different microscope position. C. Spike rates generated by S2 treatments of increasing [Ca^2+^]_ext._. Error bars = mean ± SD, n = 3. ** = p≤ 0.01, *** = p ≤ 0.001. D. Ca^2+^-GCaMP output was imaged at 146 ms. Spike shape was analysed with a bespoke MATLAB script that, after adjusting the contrast and correcting for photo-bleaching, calculated spike intensity as the mean of pixel intensity over each cell. E. Ca^2+^-GCaMP spike output during the switch from pH 4.5 (Stage 1) to pH 7.5 (Stage 2) in the presence of 5 mM Ca^2+^. F. The reverse experiment to that shown in E. G. Comparison of Ca^2+^-channel blockers on Ca^2+^-GCaMP spiking. From a Stage 1 Ca^2+^ concentration, at Stage 2 cells were exposed to 5 mM Ca^2+^ and the same concentrations (0.5 mM) of verapamil, diltiazem or nifedipine, or 10 mM BAPTA, during Stage 2. Ca^2+^ –GCaMP spikes/cell/min were expressed as a percentage of the spike rate of the no-blocker control. Error bars = mean ± SD. Data includes at least 2 biological replicates.

Investigation of the relationship between extracellular calcium, [Ca^2+^]_e_ and Ca^2+^-GCaMP responses showed that spiking frequency positively correlated with [Ca^2+^]_e_. Within the range of 1 – 10 mM, values increased from 0.03 to 0.2 spikes/cell/min (Fig 1C), but did not significantly increase further on exposure to 200 mM Ca^2+^. Ca^2+^-GCaMP spiking in the presence of 5 mM [Ca^2+^]_e_ was completely abolished by addition of 10 mM BAPTA, a Ca^2+^-specific chelator, indicating that spiking resulted from Ca^2+^influx through the plasma-membrane (Fig 1G). Changes in [Ca^2+^]_e_ from 1 – 200 mM [Ca^2+^]_e_ did not affect ambient Ca^2+^-GCaMP levels, suggesting that cellular Ca^2+^-homeostasis mechanisms adequately maintained cytoplasmic [Ca^2+^] under these conditions (Fig 1B).

Use of the aequorin Ca^2+^ reporter in *C. albicans* requires a pH of ∼ 7 or above (Sanglard, 2021). To test whether this was also true for GCaMP, the pH in Stage 1 was set at either 4.5 or 7.5 and the pH value was reversed during Stage 2. During Stage 1 at pH 4.5, spiking was negligible but commenced at a rate of 0.026 ± 0.019 spikes/cell/min on the switch to pH 7.5 in Stage 2 (Fig. 1E). In the reverse experiment when the Stage 1 baseline was set at pH 7.5, spike-rate became negligible on the switch to pH 4.5 at Stage 2(Fig. 1F). Therefore, as seen for aequorin, optimum activity of the GCaMP reporter was observed when the external pH was maintained at, or slightly above, neutral. Taken together with the finding in *S. cerevisiae* that exposure to alkaline pH triggers immediate Ca^2+^ uptake via Cch1 (Viladevall et al., 2004), the requirement for neutral pH for imaging Ca^2+^-flux is linked to the biology of the cell and not to a property of the reporter.

These experiments defined optimal conditions in which to characterise the immediate Ca^2+^– GCaMP response in *C. albicans* yeast cells to commonly-applied *in vitro* stress treatments. All further experiments were therefore conducted at pH 7.5 with a Stage 1 [Ca^2+^]_ext_ of 5 mM and an imaging frame rate of 5 s as these parameters adequately captured changes in spiking frequency between Stages 1 (baseline), 2 (treatment) and 3 (washout) without causing significant photo-stress.

We next examined the ability of 3 clinic-based inhibitors that have commonly been used with the intention of blocking Ca^2+^influx through the L-type Ca^2+^-channel homolog in *C. albicans,* Cch1. For comparative purposes, equimolar concentrations (0.5 mM) of the 2 non-dihydropyridines, diltiazem (a benzothiazepine) and verapamil (a phenylalkylamine), or the dihydropyridine, nifedipine, were added at Stage 2 where the [Ca^2+^]_e_ was held at 5 mM. At the concentrations used, none of the channel-blockers abolished Ca^2+^-GCaMP spiking during the first 10-min treatment but instead progressively inhibited spiking upon each following treatment (Fig 1G). Verapamil had the slowest kinetics and reduced spiking to ∼ 50% of the control by Treatment 3, ultimately achieving the same level of inhibition as diltiazem. The finding that verapamil does not completely inhibit Ca^2+^-uptake is consistent with previous findings in *S. cerevisiae* (Teng et al., 2008) and in *C. albicans* (Brand et al., 2007). Nifedipine was the most effective of the clinical channel-blockers by the end of Exposure 3, reducing spiking by 92.5 % of that in the control strain. However, none of the inhibitors was as effective at blocking Ca^2+^-GCaMP spiking as use of 10 mM BAPTA (i.e. a 2:1 ratio with [Ca^2+^]_cyt_), which completely abolished spiking immediately on first exposure.

### Equimolar osmotic shock caused cell shrinkage but Ca^2+^-GCaMP responses were ion-dependent

Ca^2+^-signalling is involved in a number of stress responses in *C. albicans*. We therefore imaged Ca^2+^-GCaMP dynamics during the application of a stress compound at Stage 2 for 10 min, followed by wash-out (Stage 3). The exposure was repeated at 2 further microscope positions to look for stress adaptation. To test osmotic stress, cells were exposed to equivalent ionic concentrations (2 osmol/L) of either 1 M NaCl or 0.666 M CaCl_2_. During the first exposure to 1 M NaCl, cell volume shrank by ∼ 22 %, as observed previously (Ene et al., 2012), and Ca^2+^-GCaMP6 spiking almost ceased (Fig. 2A,B,C, Movie S2). On commencement of Stage 3 (washout), cells regained their initial volume and spiking recommenced with a short period (∼ 2 min) of intense spiking before resumption of a steady rate. This was accompanied by a transient rise in ambient Ca^2+^-GCaMP, which was not seen in the empty-vector control. Similar results were seen during the 2 further exposures to 1 M NaCl, suggesting that there was an intense influx of Ca^2+^ on removal of high [NaCl]. Imaging of the empty-vector control showed that 1 M NaCl produced a slight increase in autofluorescence during the treatment phase, but this disappeared immediately on commencement of Stage 3. Despite the loss of cell volume, viability was unaffected by this short period of osmotic shock as Ca^2+^-spiking recommenced and cells were PI-negative after 3 treatments (Fig. S4A).

**Fig. 2:**
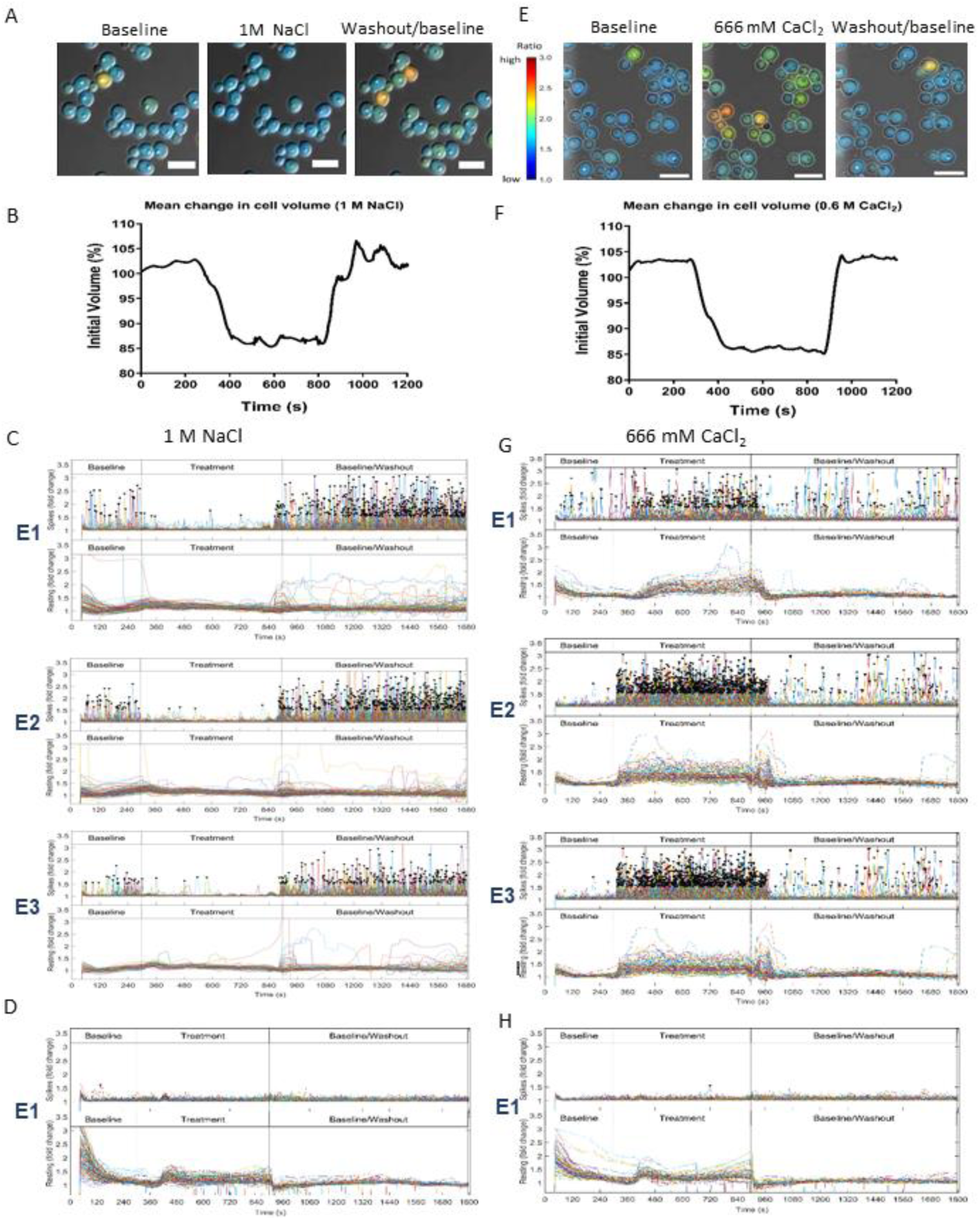
Equi-molar osmotic shocks caused cell shrinkage and differential ion-dependent Ca^2+^-GCaMP responses. A. Ca^2+^-GCaMP imaging in response to 1NaCl. B. Effect of 1 M NaCl on cell volume. The volume of all cells per field of view at each time point was normalised to the initial volume (n = 3). C. Exposure to 1 M NaCl shuts down GCaMP Ca^2+^ spiking. E1-E3 = exposures 1-3. Top panels: Ca^2+^-GCaMP spike output. Bottom panels: Ambient signal. D. No Ca^2+^-GCaMP spiking was seen in the empty-vector (A569) control strain but a slight rise in ambient fluorescence during exposure to 1 M NaCl. E. Ca^2+^-GCaMP imaging in response to 0.66 M CaCl_2._ F. Effect of 0.66 M CaCl_2_ on cell volume was determined as described. G. Exposure to 0.66 M CaCl_2_ Ca^2+^-GCaMP increased spike rate (top panels) and caused a slight rise in ambient fluorescence. H. No Ca^2+^-GCaMP spiking was seen in the empty-vector control strain but a slight rise in ambient fluorescence in the presence of 0.66 M CaCl_2_. Size bars = 10 µm.

When the experiment was repeated using 0.666 M CaCl_2_ as an alternative and equimolar salt stress, cell volume decreased by ∼ 17 %, indicating that shrinkage is a general response to osmotic stress (Fig. 2E,F, Movie S3). However, in contrast to the spiking shut-down seen in response to NaCl, the spike rate increased 6.3-fold over baseline in the presence of high [CaCl_2_]_ext_, indicating that Ca^2+^ channels retained function during cell shrinkage. Loss of Ca^2+^-GCaMP6 spiking in the presence of NaCl was therefore likely to be due to the high external Na^+^:Ca^2+^ ratio in which Ca^2+^ entry was blocked and is consistent with the short, population-wide rise in ambient Ca^2+^-GCaMP signal at commencement of NaCl washout. Exposure to 0.666 mM CaCl_2_ elicited the same level of autofluorescence in the empty-vector control as observed for 1 M NaCl (Fig. 2H), suggesting that autofluorescence and cell shrinkage are general cell responses to osmotic stress and are not ion-specific.

### Exposure to SDS elicits a 3-phase Ca^2+^ signature accompanied by major cell death and adaptation in the surviving population

The anionic surfactant, sodium dodecylsulfate (SDS), de-stabilises plasma-membrane integrity, which activates the Ca^2+^-calcineurin signalling pathway (Cruz et al., 2002). In our system, the first exposure to SDS (0.05 %) caused an intense but short-lived, 40 – 60 s Ca^2+^-GCaMP spike-burst (Fig. 3A, Movie S4). This was immediately followed by a rise in ambient Ca^2+^-GCaMP, which was not seen in the empty-vector control (Fig. 3C). The rise could be due to unregulated Ca^2+^-influx across the compromised plasma-membrane. There was no further spiking after the initial burst and ambient Ca^2+^-GCaMP levels gradually declined during the 10-min treatment, suggesting the activity of cytosolic Ca^2+^ homeostasis regulators. During the subsequent two exposures to SDS, the initial spike-burst reduced in intensity and lengthened in duration, with no reappearance of the initial rise in ambient Ca^2+^-GCaMP activity. After the initial first shock, the cell response appeared to be one of gradual recovery on subsequent treatments with SDS.

As ambient Ca^2+^-GCaMP fell during the first treatment, a large number of cells underwent a significant long-term rise in autofluorescence, which was also seen in the empty-vector control. The onset of autofluorescence occurred in heterogeneous population of cells and in an unsynchronised manner on each SDS treatment. Propidium iodide was flowed into the system after Exposure 3 and the fluorescent output of individual cells at each microscope position were compared to their end-point viability status. This revealed that only 20 % of the cells that exhibited spiking activity prior to the first exposure to SDS survived all 3 exposures, and the remaining 80 % of the population were the high-autofluorescence cells and were PI positive at the end of the experiment (Fig 3E). Cell death was dependent on the presence of extracellular [Ca^2+^], as the addition of BAPTA during SDS treatment rescued cell viability. However, the number of spiking cells that underwent cell death decreased at each exposure and 98 % of cells still spiking after Exposure 2 survived the final treatment. The gradual reduction in the severity of the spike-burst, the ambient Ca^2+^-GCaMP rise and heterogeneous autofluorescence across the 3 exposures to SDS suggest that, while the majority of the cell population did not survive the first exposure, those that did were either pre-resistant or underwent an adaptation process that promoted resistance. However, while spiking was indicative of cell viability, spiking activity was not, in itself, essential for cell survival in SDS as 7 % of non-spiking cells were nevertheless viable after Exposure 3. The phosphatase, calcineurin, is required for plasma-membrane integrity, so we next compared these responses with those of the *cnb1*Δ null mutant. No spiking was observed at any point in the experiment, even during the Stage1 pre-SDS-treatment, indicating that calcineurin is required for Ca^2+^-GCaMP spiking (Fig. 3D). Large numbers of cells exhibited autofluorescence on Exposures 1 and 2 but this was not observed during Exposure 3. All the cells were PI positive after 3 exposures to SDS so the failure of cells to autofluorescence during Exposure 3 strongly suggests that the entire population had lost viability during Exposures 1 and 2 (Fig. 3E).

**Fig. 3:**
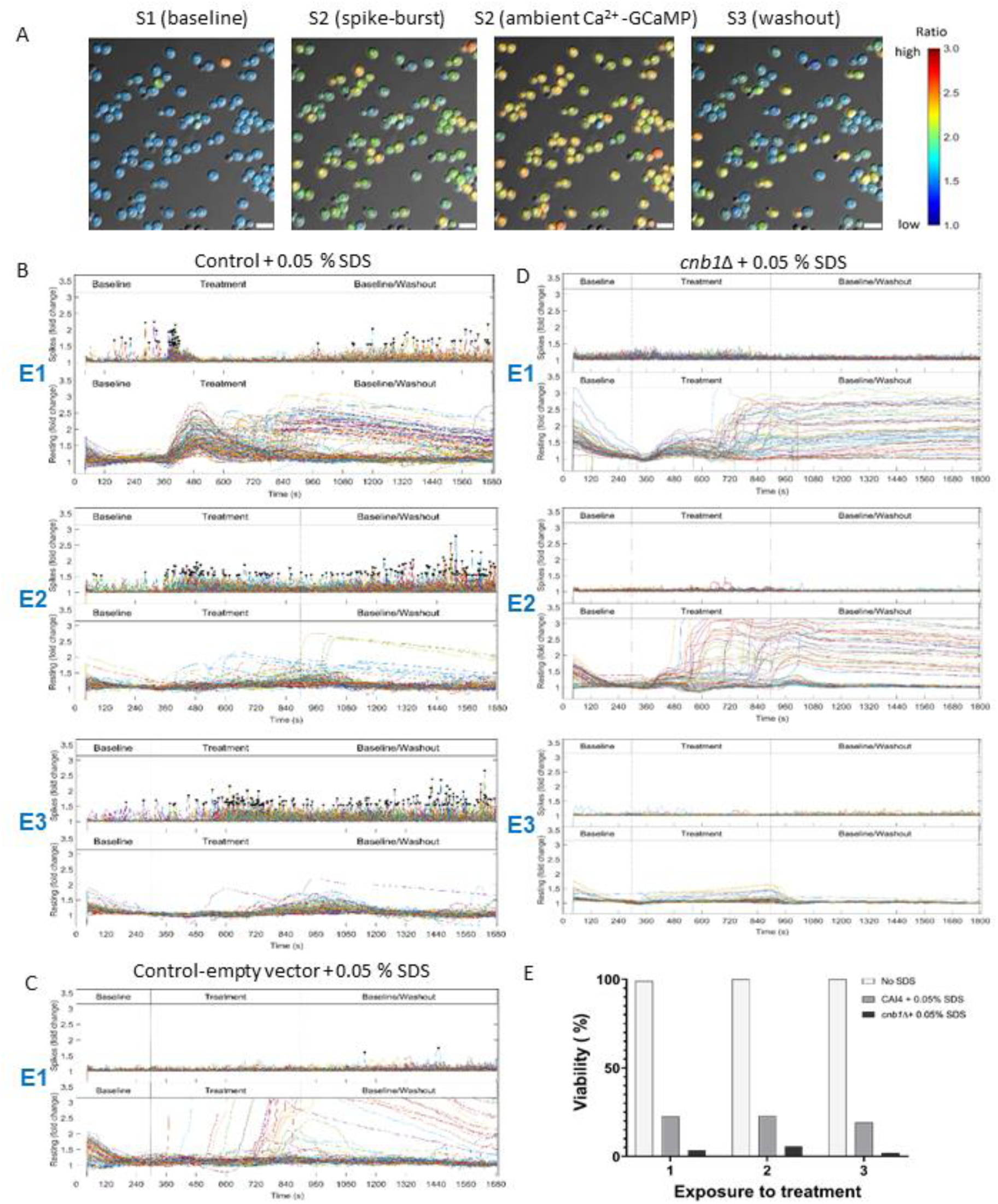
Exposure to SDS causes a 3-phase response associated with significant cell death but surviving cells undergo calcineurin-dependent adaptation upon repeat exposure. A. Ca^2+^-GCaMP imaging revealed a 3-phase response to 0.05 % SDS. B. Imaging output plots across 3 exposures to SDS. E1-E3 = Exposures 1 – 3. C. Fluorescence output of the empty-vector strain when treated with 0.05% SDS as above. D. Ca^2+^-GCaMP imaging of the *cnb1*Δ-GCaMP strain on exposure to 0.05 % SDS. E. Effect on cell viability of exposure to SDS in the control strain and *cnb1*Δ strains expressing GCaMP. Cells were stained with 1 ug/ml propidium iodide at the end of E3 and % viability expressed as the number of PI positive cells as a % of the total number of cells at each position against BWP17 –GCAMP that was not exposed to a treatment as a control. Scale bars = 10 µm.

### *C. albicans* adaptation to repeated treatment with 5 mM H_2_O_2_

1. *C. albicans* is exposed to reactive oxygen species (ROS) in the phagolysosome of immune cells (Martínez-Esparza et al., 2009, Gazendam et al., 2014) so we examined the Ca^2+^-GCaMP response to oxidative stress using 5 mM H_2_O_2_. In our system, the first exposure to H_2_O_2_ inhibited Ca^2+^-GCaMP spiking within ∼ 1 min and was accompanied by a population-wide rise in autofluorescence, which was also observed in the empty-vector control, for the duration of the treatment (Fig. 4A,B,C, Movie S5, S6). On removal of H_2_O_2_, spiking re-commenced and the autofluorescence declined. During the 15-minute washout phase (Stage 3), a small proportion of cells underwent unsynchronised short-term increases in signal that was not seen in the empty-vector control, suggesting that these cells had raised ambient [Ca^2+^]_cyt_. However, unlike the high cell death seen on SDS treatment, the population remained viable throughout the 3 exposures to H_2_O_2_ (Fig. 4E). Inhibition of Ca^2+^-GCaMP spiking was only partial during the second exposure to H_2_O_2_, and not seen on the third, with Ca^2+^-GCaMP activity continuing throughout. Furthermore, the autofluorescence rises seen in both GCaMP-expressing cells and the empty-vector control also faded with successive treatments. This suggests that Ca^2+^-spiking and the intracellular chemistry responsible for autofluorescence were modified by an adaptation response during the 90 min time-course of the experiment, such that H_2_O_2_ no longer elicited a shock response. In support of this, we confirmed that the Cap1 transcription factor, a key driver of the Oxidative Stress Response (OSR) in *C. albicans* (Wang et al., 2006), translocated from the cytoplasm to the nucleus in 55 % of cells within 2 min and 78 % of cells within 7 min of exposure to H_2_O_2_ in our experimental system (Fig. 4F), supporting the idea that amelioration of the initial shock response was due to adaptation driven by gene-expression.

**Figure 4:**
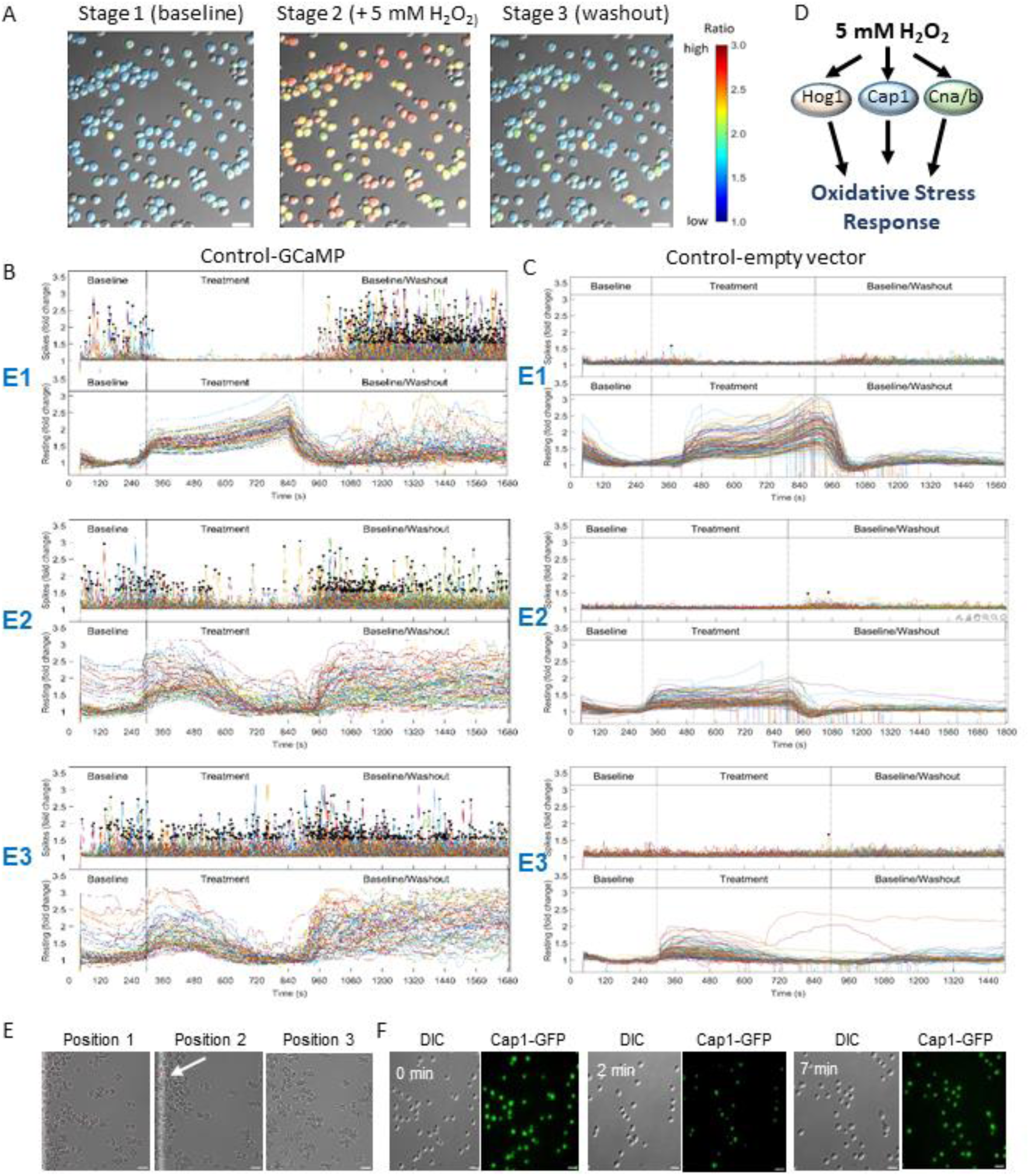
The inhibition of Ca^2+^-GCaMP spiking on exposure to 5 mM H_2_O_2_ is restored during repeated treatments. A. Ca^2+^-GCaMP imaging in wild-type cells during exposure to 5 mM H_2_O_2._ B. Imaging output plots across 3 exposures to H_2_O_2._ Top panels = Ca^2+^-GCaMP spiking. Bottom panels = ambient fluorescence. E1-E3 = Exposures 1 – 3. C. Fluorescence outputs of empty-vector strain when treated with 5 mM H_2_O_2_. D. Putative signalling pathways involved in the oxidative stress response in *C. albicans*. E. Exposure to 5 mM H_2_O_2_ does not affect cell viability. Cell populations of BWP17-GCaMP were treated with 5 mM H_2_O_2_ as before and stained with 1 µg/ml PI after exposure 3. Only one PI-positive cell was observed (arrow). F. Cap1-GFP translocation into the nucleus was imaged every 60 s after exposure to 5 mM H_2_O_2._ Scale bars = 10 µm.

### Adaptation to H_2_O_2_ requires the Cap1 transcription factor, but *hog1*Δ cells appear pre-adapted

The OSR in *C. albicans* is mediated by 2 signalling pathways, mediated by the Cap1 transcription factor and the Hog1 kinase (Fig. 4D) (Alonso-Monge et al., 2003, Enjalbert et al., 2006, Kusch et al., 2007, Zhang et al., 2000). We examined whether these pathways were required for the restoration of Ca^2+^ influx during exposure to H_2_O_2_ by expressing GCaMP6 in the respective gene deletion mutants. The first exposure of *cap1*Δ-GCaMP to H_2_O_2_ yielded very similar responses in Ca^2+^-GCaMP spiking and autofluorescence to those seen in the wild-type control cells – a shut-down in spiking and the generation of autofluorescence (Fig. 5A, Movie S7). However, the subsequent amelioration of these responses observed in control strain during Treatments 2 and 3 was absent in the *cap1*Δ mutant, indicating that Cap1 was required for cell adaptation that permitted continued Ca^2+^-GCaMP activity during oxidative stress.

**Fig. 5:**
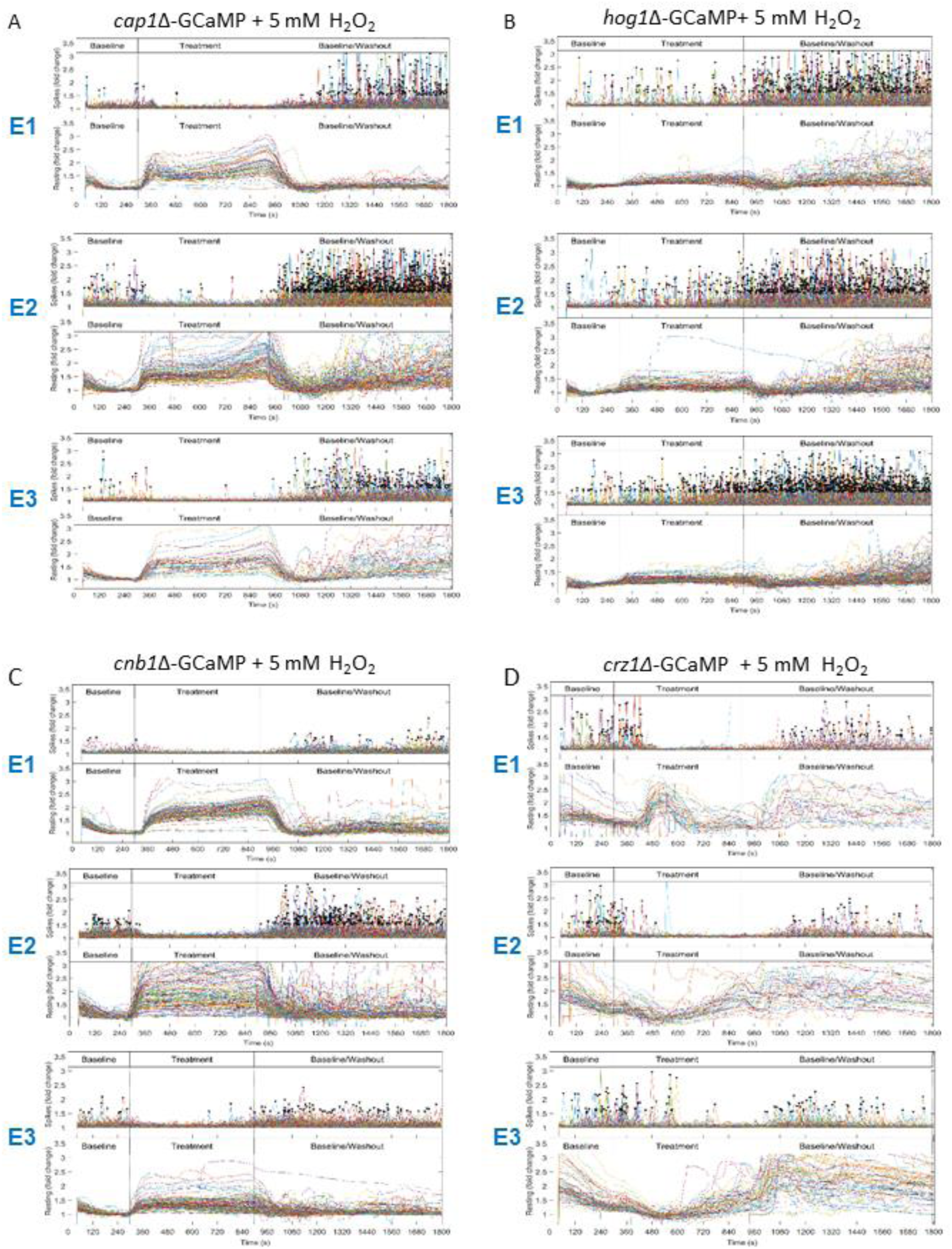
Restoration of Ca^2+^-GCaMP spiking in 5 mM H_2_O_2_ requires Cap1 and calcineurin and is only partially dependent on the Crz1 transcription factor. Spiking was not inhibited in *hog1*Δ. Ca^2+^ GCaMP spike (top panels) distributions and ambient fluorescence (bottom panels) of cells exposed to 5 mM H_2_O_2_, across the 3 stage time course during Exposures 1-3 (E1-E3). A. *cap1Δ*-GCaMP B. *hog1Δ*-GCaMP C. *cnb1Δ*-GCaMP D. *crz1Δ*-GCaMP.

The MAPK, Hog1 (High Osmolarity Glycerol 1), is phosphorylated and accumulates in the nucleus during oxidative stress, where it confers resistance to high levels of ROS (Alonso-Monge et al., 2003, Smith et al., 2004), although its effectors do not appear to include transcription factors (Dantas et al., 2015). Remarkably, in our system, unlike the shut-off of Ca^2+^-GCaMP activity seen in control cells during Treatment 1, the *hog1*Δ/GCaMP strain was minimally affected as spiking continued and autofluorescence rose only slightly (Fig. 5B, Movie S8). Treatments 2 and 3 elicited increasingly milder responses. The slight shifts seen on wash-out (an increase in spike-rate and decrease in autofluorescence) indicate that cells were able to sense the treatment but the overall response profile was very similar to that seen during the ‘adapted’ response in the control cells. This suggests that the molecular mechanisms required to maintain Ca^2+^ –spiking and suppress large rises in autofluorescence during oxidative stress were pre-initialised in the *hog1*Δ/GCaMP mutant.

### Adaptation to oxidative stress requires calcineurin but is only partially dependent on the transcription factor, Crz1

A third pathway proposed to be involved in the response to oxidative stress is the phosphatase, calcineurin. Calcineurin function has been best characterised in terms of its role in combatting membrane and Ca^2+^-stress, and its activation of gene expression via the Crz1 transcription factor (Blankenship and Heitman, 2005, Cruz et al., 2002, Karababa et al., 2006). We examined the importance of calcineurin to the OSR by expressing GCaMP6 in cells lacking the regulatory subunit, *cnb1*Δ. During Treatment 1, the response signature to H_2_O_2_ was very similar to that seen in *cap1*Δ, where spiking ceased and autofluorescence remained elevated throughout the treatment period (Fig. 5C, Movie S9). As in the *cap1*Δ mutant, there was no major change in the Ca^2+^-GCaMP6 or autofluorescence signature at Treatments 2 and 3, suggesting that cells were unable to adapt to H_2_O_2_ and that calcineurin therefore plays a role of equal importance to that of Cap1 in restoring Ca^2+^-flux and reducing autofluorescence in the presence of oxidative stress.

To further probe the contribution of calcineurin-associated effectors to the OSR, GCaMP was expressed in the *crz1*Δ null mutant. In *crz1*Δ, the Ca^2+^-GCaMP6 signature on Treatment 1 was somewhat erratic but essentially the same as that seen in control cells, with a cessation in spiking and a rise in autofluorescence (Fig. 5D, Movie S10). However, unlike the *cnb1*Δ mutant, Ca^2+^ spiking resumed and autofluorescence reduced during subsequent treatments. In addition, as in the control strain, there was a rise in ambient Ca^2+^-GCaMP6 during the wash-out phase. Therefore, even though ambient Ca^2+^-GCaMP6 activity became increasingly disorganised and heterogeneous, it appeared that the *crz1*Δ mutant was able to undergo adaptation to oxidative stress. Crz1 is responsible for transcription of 69 calcineurin-dependent genes, 5 of which (including Golgi Pmr1 and vacuolar Pmc1) are required for Ca^2+^ transport and homeostasis (Karababa et al., 2006, Xu et al., 2019) so it is possible that the erratic ambient Ca^2+^ signature seen on Crz1 deletion was due to the cells’ inability to regulate homeostasis during onset of the stress response. The finding that calcineurin, but not Crz1, is required for adaptation points to the importance of alternative calcineurin effectors for delivering a normal response to oxidative stress.

### The plasma-membrane Ca^2+^-channel, Cch1, and extracellular Ca^2+^, but not the vacuolar TRP-channel, Yvc1, is required for adaption to oxidative stress

Influx of Ca^2+^ into the cytoplasm requires entry through the plasma-membrane from the extracellular environment or release from intracellular stores. Currently, the only known channels that mediate cytoplasmic Ca^2+^ inflow are the plasma-membrane-localised, Cch1, which is most closely related to non-voltage-gated sodium leak channels (NALCN), and Yvc1, a TRP channel that releases Ca^2+^ from the vacuole (Liebeskind et al., 2012, Palmer et al., 2001). Interestingly, in resting *cch1*Δ cells, weak Ca^2+^-GCaMP spiking was seen during Stage 1 and during the Stage 3 washout period (Fig. 6A, Movie S11), suggesting the existence of an alternative channel for Ca^2+^-entry through the plasma-membrane and/or that Ca^2+^ was released from intracellular stores. During exposure to H_2_O_2_, however, the spike-rate was greatly reduced during each exposure, suggesting no restoration of Ca^2+^-influx in this mutant. Conversely, the high level of autofluorescence seen on Exposure 1 was gradually reduced to negligible by Exposure 3 and the heterogeneous rises in ambient Ca^2+^-GCaMP seen in the control strain were muted throughout. In the *yvc1*Δ null mutant, the response and adaptation to repeated H_2_O_2_ exposure was very similar to that seen in the control strain (Fig. 6B, Movie S12). However, when extracellular Ca^2+^was chelated by BAPTA, spiking was completely abolished and the perturbation in autofluorescence in the *yvc1*Δ null was minimal (Fig 6C), so Ca^2+^ release from the vacuole via Yvc1 does not contribute to Ca^2+^-GCaMP spiking but may alter the intracellular chemistry that generates autofluorescence during oxidative stress. Taken together, these results suggest that there is a low-conductance plasma-membrane channel in addition to high-conductance Cch1 in *C. albicans*, and is consistent with the proposed existence of a Low-Affinity Calcium System (LACS) and High-Affinity Channel System (HACS) (Fischer et al., 1997, Paidhungat and Garrett, 1997, Iida et al., 1990, Muller et al., 2001). This low-conductance channel is unable to adapt from its partial inhibition by H_2_O_2_ and neither channel plays a role in the rise and adaptive recovery of intracellular autofluorescence on repeated treatments, which occurred in both *cch1*Δ and *yvc1*Δ mutants, unlike the *cap1*Δ and *cnb1*Δ strains.

**Fig. 6:**
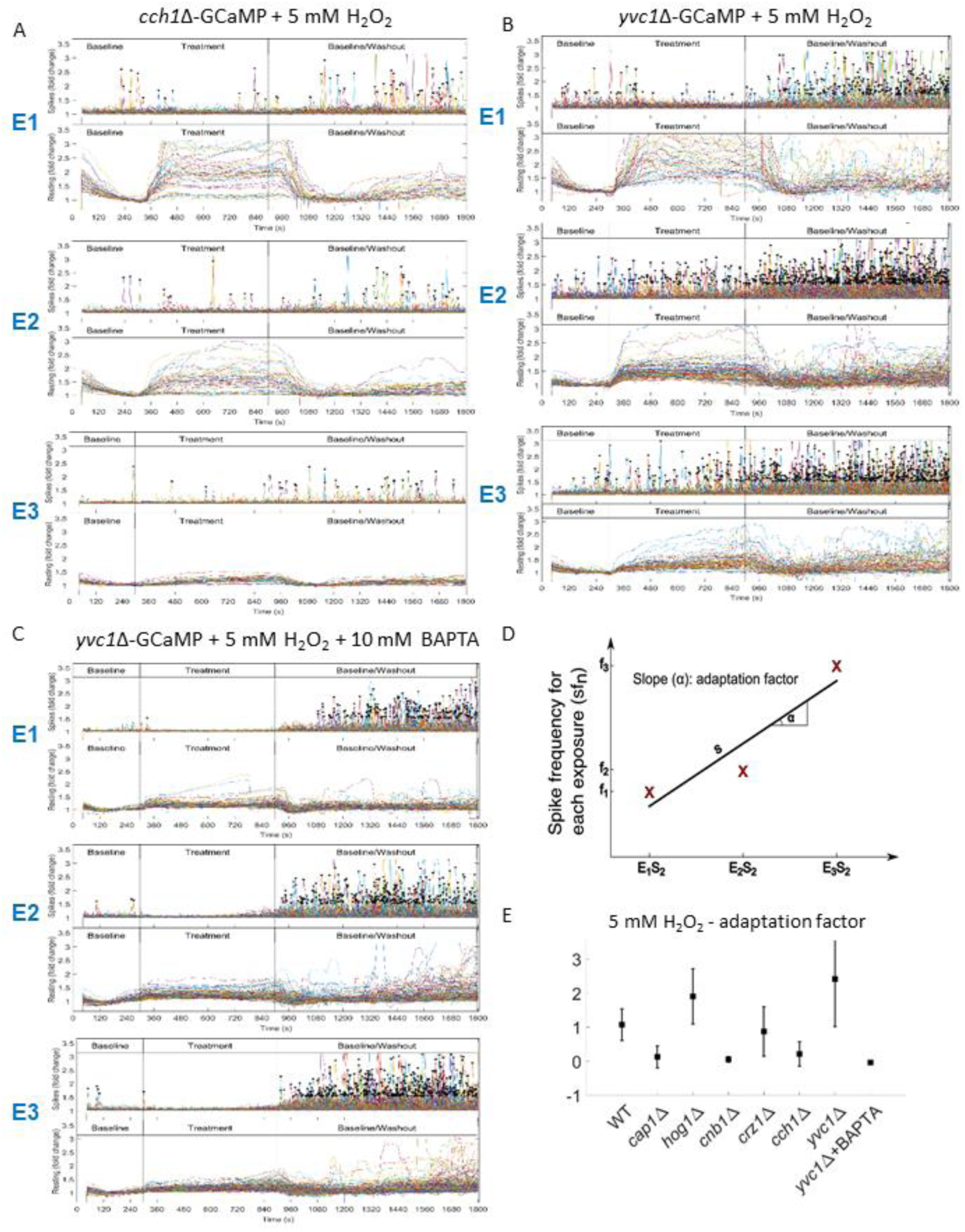
Cch1 and extracellular Ca2+, but not Yvc1, are required for adaptation to H_2_O_2_ treatment. Ca^2+^ GCaMP spike (top panels) distributions and ambient fluorescence (bottom panels) of cells exposed to 5 mM H_2_O_2_, across the 3 stage time course during Exposures 1-3 (E1-E3). A. *cch1Δ*-GCaMP B. *yvc1Δ*-GCaMP C. *yvc1Δ*-GCaMP + 10 mM BAPTA. D. Definition of the adaptation factor (AF) as the change in spike frequency per cell from exposure to exposure. E. Adaptation factors of wild-type and mutant strains on treatment with 5 mM H_2_O_2_. Student’s t tests (with Bonferroni correction for multiple comparisons) show that the wild type, *hog1*Δ, *crz1*Δ and *yvc1*Δ adapt to H_2_O_2_ (with AF significantly different to 0), whereas *cap1Δ*, *cnb1Δ*, *cch1Δ* and *yvc1Δ*+BAPTA do not (AF not significantly different to 0).

### Restoration of Ca^2+^-GCaMP spiking on repeated exposure to H_2_O_2_ in signalling pathway mutants

In order to compare adaptation responses in the wild-type strain and signalling pathway mutants to repeated H_2_O_2_ exposure, the mean Ca^2+^-GCaMP spike rate was plotted across the 3 exposures to H_2_O_2_ for each strain (Fig. 6D,E, S5) and the Adaptation Factor determined by linear regression. The *cap1*Δ, *cnb1*Δ and *cch1*Δ mutants showed little or no restoration of Ca^2+^-GCaMP spiking after the shut-off seen during the first exposure to H_2_O_2_, unlike the wild-type control strain and the *crz1*Δ and *yvc1*Δ mutants (Fig. 6E). The *hog1*Δ mutant also adapted but from a different starting position compared to the control strain, as spiking in this mutant was not inhibited on the first exposure H_2_O_2_. However, strains that underwent adaptation to H_2_O_2_ were differentially affected in other ways, for example, autofluorescence commenced at a high level in *yvc1*Δ but reduced over time, whilst in the *hog1*Δ mutant the autofluorescence level was only slightly raised throughout. Therefore, Ca^2+^-GCaMP spiking behaviour is a valuable indicator of cell viability and active homeostasis mechanisms but the other fluorescent signals provide additional insights into physiological aspects of cell responses.

### Exposure to the antifungal drugs, Caspofungin and Fluconazole, does not elicit an immediate Ca^2+^ stress response

Cell systems that confer resistance to antifungal drugs are of great interest in medical mycology. We therefore used our system to test whether exposure to fluconazole (8 µg/ml), which targets ergosterol biosynthesis, or Caspofungin (0.032 µg/ml), which inhibits cell-wall biosynthesis, elicit changes in Ca^2+^-GCaMP6 activity. Cells were exposed to two 10 min treatments but no change in Ca^2+^-GCaMP6 activity or ambient Ca^2+^-GCaMP6 levels were observed (Fig. S2). The lack of an immediate response to these antifungal compounds indicate that they are not sensed by cells in such a short time-frame.

## Discussion

### The use of genetically-encoded Ca^2+^ reporters in fungi

Despite the importance of Ca^2+^-signalling and homeostasis in fungal pathogens, our understanding of real-time Ca^2+^-dynamics in single cells has been limited by the paucity of effective reporters in fungal cells. The development of new genetically-encoded [Ca^2+^] indicators (GECIs) has allowed examination the role of Ca^2+^ flux in the growth of filamentous fungi. The Cameleon FRET (Förster Resonance Energy Transfer) ratiometric system used Ca^2+^-binding to Calmodulin and the M13 calmodulin-binding peptide to transfer energy from CFP to YFP to visualise cytosolic Ca^2+^ (Kim et al., 2012, Miyawaki et al., 1997). The Cameleon reporter was expressed in the filamentous fungi, *Fusarium oxysporum, F. graminearum* and *Magnaporthe oryzae,* where real-time imaging revealed pulsatile changes of ∼ 26 s in duration in apical Ca^2+^ during hyphal growth for the first time. Tip-focussed Ca^2+^ spiking was observed in *Colletrotrichum graminicola* using the same reporter, although spiking did not correlate with the non-pulsatile tip growth rate. More recently, Ca^2+^-reporters have been developed from GCaMP, a construct using the Ca^2+^-Calmodulin-M13 peptide interaction to excite circularly-permutated GFP (cpGFP) (Tian et al., 2009, Takeshita et al., 2017, Nakai et al., 2001). R-GECO was a GECI derived from GCaMP3, the third iteration of GCaMP, and its visualisation *A. nidulans* and *N. crassa* led to a model of oscillatory Ca^2+^-entry followed by actin disassembly and vesicle exocytosis, a cycle that correlated with the step-wise nature of hyphal tip extension in these fungi (Takeshita et al., 2017). GCaMP6 has been adapted for expression in *Aspergillus fumigatus*, where heterogeneous Ca^2+^ waves were imaged in hyphae and germlings and dynamics were altered by exposure to high (200 mM) Ca^2+^ (Muñoz et al., 2021). It has also been used in *Schizosaccharomyces pombe*, where transients during cell-separation events, hypo-osmotic and Ca^2+^ shock were observed (Poddar et al., 2021). Here, we engineered a codon-optimised version of GCaMP6f in order to visualise Ca^2+^-dynamics in the yeast form of the fungal pathogen, *C. albicans*, for the first time and dissect their role in stress responses.

## Analysis and model for Ca^2+^-GCaMP6 outputs in *C. albicans*

The imaging output from our perfused microfluidics system showed 3 variants of fluorescence output from GCaMP-expressing cells, two of which split further on comparison of output from the empty-vector control strain (Fig 7A).

**Fig. 7:**
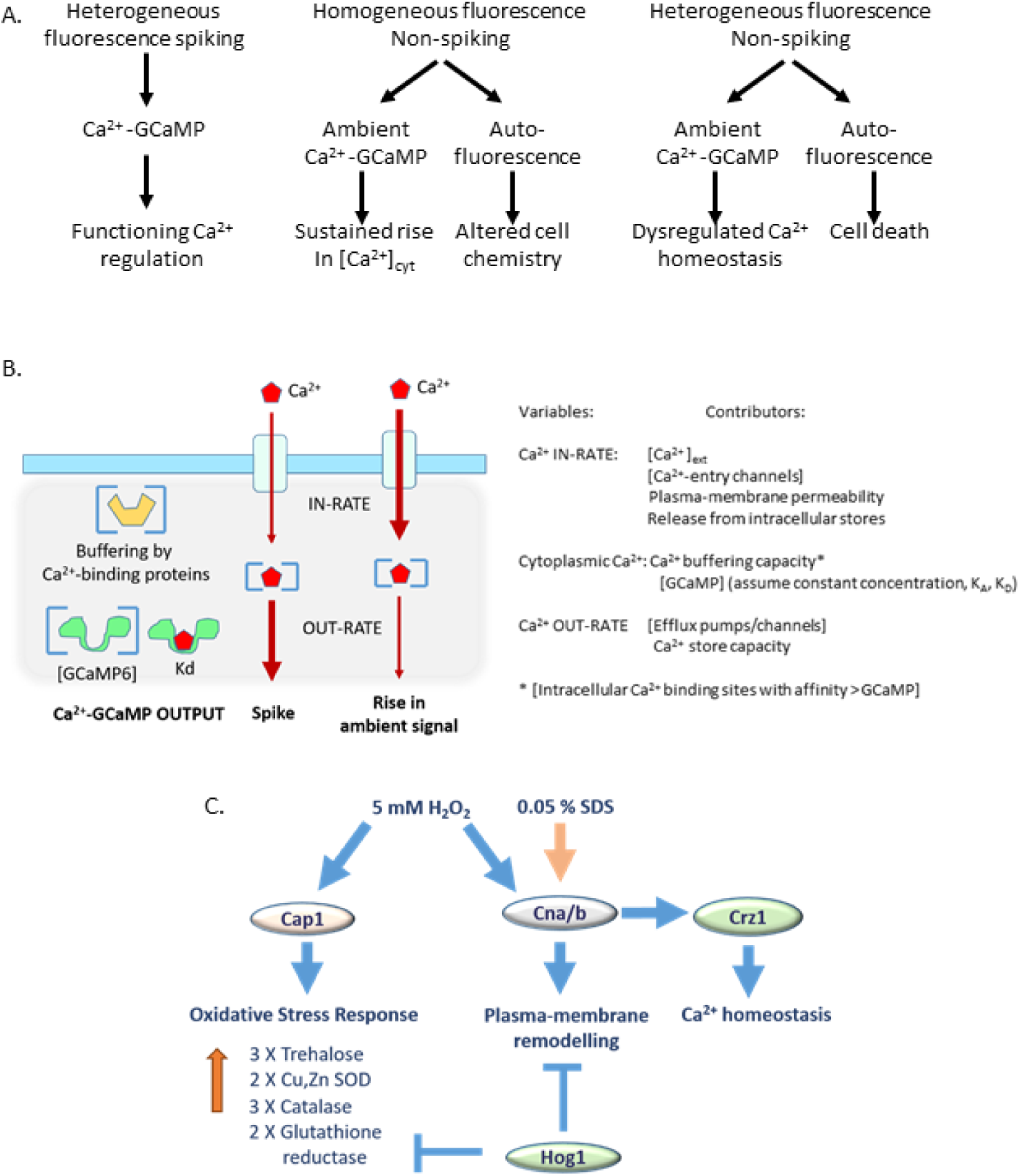
A. Categorisation of fluorescence outputs from GCaMP imaging in *C. albicans*. See text for details. B. Proposed mechanisms for how Ca^2+^ influx and efflux rates are affected by variables but together deliver Ca^2+^ GCaMP spiking vs changes in ambient Ca^2+^ GCaMP levels. C. Model for adaptation to oxidative and membrane stress in *C. albicans*. Oxidative stress requires Cap1 and calcineurin signalling for upregulation of antioxidant enzymes and remodelling of the plasma-membrane. Ca^2+^ GCaMP spiking is not inhibited in the *hog1Δ* mutant as it is pre-adapted through release of gene-expression inhibition, which may reduce plasma-membrane permeability to H_2_O_2_. Calcineurin is necessary for adaptation and responses to both H_2_O_2_ and SDS, also via plasma-membrane remodelling. Crz1 is required for maintenance of Ca^2+^ homeostasis during stress responses.

1. *Heterogeneous Ca^2+^-GCaMP spiking in individual cells* was unsynchronised in both resting cells and on application of a stress treatment. Therefore, this heterogeneity suggests that the ability of a cell to produce a Ca^2+^ spike may depend on an intrinsic cell state. Cells undergoing Ca^2+^-GCaMP spiking were assumed to be viable with a functioning Ca^2+^-regulatory system as cells that stopped spiking, for example in the presence of 1 M NaCl, were able to re-start on removal of the stress. Imaging in the millisecond range showed that spike-duration was 5 – 6 s, with a time-to-peak of 1 – 2 min, much slower than the tens-of-milliseconds for GCMP6f in neuronal cells (Chen et al., 2013). However, this fast frame-rate could not be sustained for the 30 min period required for each stress exposure due to photo-stress. Imaging every 5 s satisfactorily captured spike rates whilst minimising photo-stress during the period of interest (30 min) (Fig. S2). However, this rate did not allow examination of inter-spike intervals in single cells, which are predicted to have an interval minimum that may vary according to channel number, regulation and environmental conditions.
2. *Homogeneous non-spiking intracellular fluorescence* across the population occurred in response to exposure to high [Ca^2+^]_ext_, H_2_O_2_, the first treatment with SDS and on washout of 1 M NaCl. While the signal intensity varied by cell, these changes followed the same overall magnitude and time-course trend across the cell population. Comparison of the fluorescence output between cells expressing GCaMP vs an empty vector distinguished between Ca^2+^-GCaMP output, which we termed ‘ambient Ca^2+^-GCaMP’, and autofluorescence. Ambient Ca^2+^-GCaMP was raised markedly during the first few minutes of exposure to SDS and also slightly at the commencement of washout of 1 M NaCl, while whole-population autofluorescence was seen during exposure to H_2_O_2_. As none of these whole-population responses induced loss of cell viability (Fig. S4), we concluded that rises in ambient Ca^2+^-GCaMP were due to a sustained rise in ambient [Ca^2^]_cyt_ (see model below), while rises in autofluorescence during treatment phases were due to altered intracellular chemistry, particularly during exposure to H_2_O_2_.
3. *Random incidence of fluorescence in a heterogeneous sub-population* of cells was seen on treatment with SDS or H_2_O_2,_. In SDS, this was due to autofluorescence and occurred in a large number of cells, which commenced in an unsynchronised manner during the latter period of exposure and was sustained in individual cells through the washout stage. Propidium iodide staining after the 3^rd^ exposure to SDS and tracking the output from each PI-positive cell back across the time-course, showed a positive correlation between cell autofluorescence and its subsequent death. The sub-population of cells that did not autofluoresce survived all 3 treatments with SDS. The autofluorescence signal in the *cnb1*Δ mutant was additionally informative because it revealed that all cells had died on the second exposure to SDS (Fig. 3D). In contrast, the heterogeneous fluorescence after H_2_O_2_ exposure was seen in a sub-population of GCaMP-expressing strains and not in the empty-vector control. The signal was therefore due to raised ambient Ca^2+^-GCaMP levels.

Although the signals rose and fell erratically in individual cells, they appeared to be generated by dysregulated Ca^2+^ homeostasis and did not lead to cell death, unlike the sustained autofluorescence seen in SDS.

The cellular mechanisms underpinning the generation of spiking vs ambient Ca^2+^-GCaMP signals require further exploration but may depend on the ratio of cytoplasmic [Ca^2+^] to Ca^2+^-GCaMP molecules, which in turn is defined by the IN-rate vs the OUT-rate of Ca^2+^-flux (Fig 7B). Our proposed model is that, if the OUT-rate (i.e., through the removal of cytoplasmic Ca^2+^ via plasma-membrane pumps or into intracellular stores) is greater than the IN-rate, then any Ca^2+^ ions entering the cell that that bind GCaMP will, on dissociation, be quickly removed from the cytoplasm. Conversely, if the IN-rate is greater than the OUT-rate, then [Ca^2+^]_cyt_ will increase and dissociated Ca^2+^ will remain in the cytoplasm to repeatedly re-bind to GCaMP, thus yielding a sustained rise in fluorescence. The many potential variables that may affect this balance remain to be investigated. These include buffering by indigenous Ca^2+^-binding sites, the status of intracellular stores and the function of Ca^2+^-channel regulators, all of which may be altered by changes in the intracellular biochemistry wrought by stressors and by adaptive gene expression. The effect of these variables on Ca^2+^-GCaMP dynamics remain to be explored.

### Adaptation to SDS and H_2_O_2_ is likely due to plasma-membrane remodelling

Unlike the consistent shut-down of Ca^2+^-GCaMP spiking seen in high osmotic stress, cells treated repeatedly with SDS or H_2_O_2_ (with a 20-min recovery period between each exposure) exhibited an ability to adapt, whereby the third application of the stress no longer inhibited spiking. In the first treatment with SDS, a detergent, the spike-burst was followed by a population-wide rise in ambient Ca^2+^-GCaMP levels, suggesting that partial disruption of the plasma-membrane allowed a sudden influx of Ca^2+^ into the cytoplasm, which temporarily overwhelmed the homeostasis mechanisms. Around 80 % of cells did not survive this first exposure and showed rises in autofluorescence that declined slowly over 20 min or more, consistent with leakage of cytoplasm from damaged membranes. Cell death was due to Ca^2+^ overload as inclusion of 10 mM BAPTA with SDS treatment led to almost 100 % survival (Fig S4). The remaining 20 % of cells were able to generate Ca^2+^ spikes during the 2^nd^ and 3^rd^ exposures, did not display further rises in ambient Ca^2+^-GCaMP levels and survived to the end of the experiment. The first exposure to SDS may have selected a sub-population that was already partially resistant to SDS. The finding that the *cnb1*Δ mutant was unable to survive in SDS, suggests that resistance to SDS in the presence of extracellular Ca^2+^ is mediated by calcineurin and is likely to involve properties of the plasma-membrane.

Calcineurin, and the oxidative-stress related transcription factor, Cap1, were both required for restoring Ca^2+^-GCaMP spiking and homogenous, non-lethal rises in autofluorescence in 5 mM H_2_O_2_. Cap1 activity has been well-characterised in terms of its role in up-regulating antioxidant enzymes for removal of H_2_O_2_ as a key resistance mechanism in the oxidative stress response (Zhang et al., 2000, Cao et al., 2008) The finding that the *hog1*Δ mutant was pre-adapted is consistent with its increased trehalose content and up-regulated Catalase A, which are expected to provide intracellular protection against H_2_O_2_ (Gónzalez-Párraga et al., 2010, Enjalbert et al., 2006). However, in *S. cerevisiae*, no correlation was found between the capacity to remove cytoplasmic H_2_O_2_ and resistance (Branco et al., 2004). Instead, during exponential-growth, resistance to H_2_O_2_ was conferred by a two-fold decrease in plasma-membrane permeability through remodelling (Sousa-Lopes et al., 2004, Branco et al., 2004). The downstream effector of Hog1 in the oxidative stress response is not known (Dantas et al., 2015) but this study suggests that it is involved in cross-talk with calcineurin signalling and may act by repressing genes involved in membrane remodelling that ultimately controls permeability to H_2_O_2 (_Fig. 7C).

## Methods

### Generation of GCaMP6 –expressing *C. albicans* strains

The strains used in this study are shown in Table S1. G-CaMP6f (Chen et al., 2013), was synthesized by GeneArt, Germany, incorporating modifications for *C. albicans* by circular permutation of *yEGFP3* (Cormack et al., 1997, Baird et al., 1999) and codon optimisation of the Ca^2+^ Calmodulin and M13 domains (Fig. S1). GCaMP6 was subcloned between the *Age*I and *Sph*I sites in plasmid CIp-*NATflip-pACT1-ScCyct* (Shen et al., 2005, Shahana et al., 2014) to generate plasmid B153. Plasmid B251 bearing *URA3* as the selectable marker was generated by cloning *ACT1p-GCaMP6* into the *Mlu*I and *Kpn*I sites of the CIp10 plasmid (Murad et al., 2000, Brand et al., 2004). Plasmid B250, without GCaMP6, was transformed into cells as the empty-vector control. Plasmids were linearized with *Stu*I. B153 and B250 were transformed into SC5314 and B251 was transformed into the Ura-minus strain, DSY3891, to generate strains A567, A569 and A585, respectively. Transformants were selected on solid Yeast-Bacto-Peptone-Dextrose or Sabouraud-Dextrose plates containing 200 µg/ml Nourseothricin or 0.67 % Yeast-Nitrogen Base without supplements and confirmed by restriction digest, fluorescence microscopy and PCR using primers NATR fw 5-AGCTTGTTCACCATCGGAAG-3 and NATR rv 5-TTCTGTTCCAGGTGATGCTG-3 or primers SP RPSgs_Rev 5-GGTAGTCGATATTCAGGGCC-3 and GCaMP6-1R 5-CCTTCAAACTTGACTTCAGC-3.

### Imaging of Ca^2+^-GCaMP responses in yeast

Cells were grown overnight at 30 °C in SD (0.67 % (w/v) Yeast Nitrogen Base with amino acids and ammonium sulphate) medium with shaking at 200 rpm. Approx 2 × 10^6^ cells in 200 μl were washed x3 with ddH2O and re-suspended in 75 μl Modified Soll’s Medium (MSM) (Crombie et al., 1990, Brand et al., 2007) +/-Ca^2+^, but without other added metals, before imaging at 30 °C in Y04C microfluidic plates (Millipore Merck, UK) connected to an Onix microfluidic perfusion system (CellASIC Corp., USA). Prior to cell loading, the PBS from 2 wells and the cell well was flushed through with 75 μl medium by sealing the plate and applying 2 x 6 psi for 10 s, followed by 2 min at 5 psi from each medium well. The remaining medium was removed and 300 μl medium corresponding to the two experimental conditions (control and shock) were added to each well. Pressure (3 psi) was applied to the “shock” well for 1 min to reduce the lag time between pressure application and medium entering the cell well, as determined by calibration experiments using a fluorescent dye. Cells (75 μl) were loaded into the cell well with a ceiling height of 3.5 μm where 1 – 2 pulses at 6 psi for 10 s yielded ∼100 cells/field of view. MSM was perfused at 2 psi. A DeltaVision Core microscope (Image Solutions Ltd., UK) with a CoolSNAP camera (Photometrics United Kingdom Ltd., UK), a 60× objective and a GFP filter was used for live-cell imaging. After plate loading, cells were incubated for 40 min in ‘baseline’ conditions. At t = 40 min, Stage 1 imaging in DIC and GFP commenced at Position 1 at 5 s intervals for 30 min, with an Auto-focus step every 2 min. At Stage 2, t = 45 min, cells were exposed to a stress compound for 10 min before Stage 3, a return to baseline conditions with continued imaging to t = 70 min. After a 5-min interval, imaging and treatment (Exposure 2 and 3) were repeated at Positions 2 and 3, and without treatment at Position 4 as a control to confirm long-term cell viability. Alternatively, to assess cell death following 3 treatments, propidium iodide (1 µg/ml) was perfused into the chamber and cell staining imaged and quantified. To assess imaging artefacts, cells expressing the empty plasmid were imaged and GCaMP6 cells were imaged in baseline (Stage 1) conditions of 5 mM Ca^2+^ with a switch of perfusion channels but with no treatment. Baseline conditions in all experiments were set at 5 mM Ca^2+^ and pH 7.5 (to maintain pH ≥ 7 in the presence of added reagents), unless otherwise stated. To determine GCaMP spike shape, *crz1*Δ-GCaMP was grown in MSM pH 7.5, 5 mM CaCl_2_ in an Ibidi U slide (Thistle Scientific, UK) and imaged on a Dragonfly spinning disk confocal microscope (Oxford Instruments, UK) at a frame rate of 146 ms.

### Imaging of Cap1-GFP localisation

Cap1-GFP cells were grown overnight at 30 °C in YPD (2 % mycological peptone (w/v), 2 % glucose (w/v), 1 % (w/v) yeast extract with shaking at 200 rpm. Cells were harvested, washed x3 in ddH_2_O and resuspended to ∼ 1×10^6^ cells/ml. Cells (100 µl) were added to an Ibidi U slide, allowed to adhere and incubated in MSM + 5 mM H_2_O_2_ for 15 min at 30 °C. Cells were imaged in DIC and FITC every minute before the addition of 5 mM H_2_O_2,_ and at every minute thereafter for 15 min.

### Image analysis software

Two-channel image sequences were imported into MatLab® using the BioFormats package (Linkert et al., 2010). After correction for stage drift, if necessary, image series were aligned to the first time-point by (*x,y*) translation estimated using the MatLab® *imregtform* function, and applied using the *imwarp* function. Aligned images were filtered with a 5×5 (*x,y*) pixel box average to reduce noise, sub-sampled by a factor of 3 to decrease processing time by removing redundant information after smoothing, and normalised in the range [0 1]. The average background intensity was measured from a manually-defined region-of-interest (ROI) for each time-point and subtracted.

As cells did not move during the time-series, an essentially noise-free template image was constructed using additional Gaussian smoothing in (*x,y*), with σ set to the minimum cell radius (typically 4 pixels), followed by an average projection of the time-series in *t*. The initial segmentation used a mid-grey local adaptive filter, which uses a spatially varying threshold set as the mean after erosion and dilation with circular neighbourhood set by the maximum cell radius (typically 25 pixels). Touching cells were separated using a watershed transform of the Euclidean Distance Transform (EDT) of the binarised cells, with additional suppression of maxima less than 15 % of the maximum to avoid over-segmentation. Within each cell territory, the actual cell boundary was defined as 50 % of the local intensity maximum in each territory. Cells touching the border or below a minimum area (typically ∼40 pixels set by the minimum cell radius) were excluded from further analysis. Each ROI was given a unique label.

A 50 s photobleaching period was observed within the first 10 frames of fluorescence imaging and were excluded from analyses. As the signal from each cell declined due to photobleaching, the baseline fluorescence (*F_b_*) was estimated using a rolling median filter with a large window size (typically 60 frames or 5 min) to exclude spikes or transients. In cases where there was an extended shift in the saturation Ca^2+^ level, eg during H_2_O_2_ treatment, the baseline was calculated using the MatLab® *msbackadj* function, which finds points from the lowest 10 % quantile in multiple shifted windows (typically 60–120 frame or 5–10 min duration), with a typical step size of 30 frames (150 s). Local baseline values were then fit using loess (quadratic) regression.

The *δF/F_b_* ratio, and the fold change in fluorescence *F/F_b_* were calculated from the background-subtracted signal and the baseline. Spikes and changes in resting level were separated using a rolling median filter with a typical window size of 7 frames (35s). The first 10-20 frames (50-100 s) were excluded to avoid the initial peak in signal following perturbation during experiment set-up. Changes in ambient level were calculated as the fold change above baseline, after the median filter. Spikes were identified from the ratio between the background-subtracted value and the median filtered data using the MatLab® *findpeaks* function. Spikes were defined as a signal fold change of > 1.5, a prominence (difference to the next peak) > 0.2, and the maximum width < 10 frames (50 s). The number of spikes/cell was determined, along with the fold-change and width (dependent on the sampling rate, during each stage of the experiment. If there was more than one spike in a cell, the inter-spike interval was determined. The overall spike frequency was calculated as spikes cell^-1^ min^-1^ to normalise between different experiments with varying number of cells and treatment duration. Data were output to Excel. The background effect of photo-stress was quantified using control experiments but found to be insignificant during the first 20 min of imaging. Falsely identified cells, due to background in heterogeneities. were manually removed.

A pseudo-colour coded image of the Ca^2+^ transients was constructed by assigning the *F/F_b_* ratio value for each cell to the corresponding ROI, imposing display limits of [0.8 3], and converting to a rainbow colour scale running from blue to red. The coded ratio was then superimposed on the bright-field image. Software is freely available for download from (https://markfricker.org/) and (https://github.com/Callump427/C-albicans-CSA-software). Plots of GCaMP spikes and ambient GCaMP Ca^2+^ activity were generated using a bespoke MATLAB script, which imported processed data generated from previous analysis.

Changes in the intracellular ambient level of Ca^2+^-GCaMP6 on exposure to stress were differentiated from changes in autofluorescence by comparing the signals between GCaMP-expressing strains and strains expressing an empty-vector as a control. To determine changes in cell volume, cells were first segmented in the DIC channel using the Yeast Spotter segmentation software (Lu et al., 2019) in Python V3.6, using the PyCharm IDE (2021.2.3). Separated images were placed into an input directory, and the pre-defined weights for the neural network were downloaded from (http://hershey.csb.utoronto.ca/weights.zip) and placed into the same directory as the segmentation and opts.py files. The segmentation script was run resulting in FIJI compatible masks. The cells were assumed to be spheres and the volume was calculated from the radius and area of each segmented cell. The area of biological replicates was averaged, and normalised to the initial volume.

## Data analysis

In Excel, spike values were displayed in chronological order, allowing separation into Stages 1 – 3 to identify trends. The number of spikes/cell/min were quantified for baseline (Stage 1), 10 min of stress treatment (Stage 2) and 15 of washout (Stage 3). In Stage 2, spikes/cell/min were calculated in a 5 min window commencing 2.5 min after wash-in and ending 2.5 min before Stage 3 washout. Spike-rate generated by increasing [Ca^2+^]_ext_ were analysed using a one-way ANOVA and a post-hoc Dunnett’s multiple comparisons t-test.

The percentage of cells spiking in each field of view during these intervals was calculated. To compensate for variability between Ca^2+^-increase experiments, spiking was normalized with respect to the first 4 min of control. The first 10 frames were discarded due to rapid initial photo-bleaching. The ‘adaptation factor’ to H_2_O_2_ was determined by quantifying spikes/cell/min for each Exposure 1, 2 and 3 (using the 5 min window) and determining the gradient (adaptation factor) of the line of best fit across the exposures.

Ca^2+^-GCaMP output from Dragonfly confocal imaging were analysed with a bespoke MATLAB script. Individual cells of interest we cropped by hand and thresholded so that pixels below a minimum intensity were replaced with black, those above a maximum intensity were replaced with white, and linear interpolation was used for pixels with intermediated intensities. Photo-bleaching was corrected by multiplying each pixel intensity by a 1+αt, where α = 3.4×10^-3^s^-1^ and t is the time in seconds. For any given time point, the Ca^2+^-GCaMP signal was defined as the mean intensity within each cell. Traces were smoothed using the MATLAB smoothdata() function, before being scaled so that all spikes lie between 0 and 1.

## Acknowledgements

AB conceived the project and wrote the manuscript. CVG conceived the experimental design. SW designed the GCaMP reporter. AM, KL, LV-M, SC and TB constructed strains and optimised imaging. MF developed the image analysis software. CVG and CP carried out the microfluidics experiments and imaging analysis. NG assisted with preparation of the manuscript. PS, SN and DMR developed and undertook the theoretical data analysis and contributed to the interpretation of the results.

## Supplementary Figures

Fig. S1 Codon optimised nucleotide sequence of GCaMP6f for expression in *C. albicans*.

Fig. S2: Onset of photo-stress occurs in Stage 3 (washout) of the experimental time-course.

Fig. S3: 10 min exposure to fluconazole or caspofungin does not affect Ca^2+^ GCaMP activity

Fig. S4: Cell viability of control strain expressing GCaMP after 3 exposures to osmotic shock, SDS, or H_2_O_2_

Fig. S5: Adaptation Factors for WT and mutant strains treated with 5 mM H_2_O_2_

## Supplementary movies

Movie S1: BWP17-GCaMP 1 mM – 200 mM Ca^2+^

Movie S2: WT-GCaMP 1M NaCl

Movie S3: BWP17-GCaMP 0.6 M Ca^2+^

Movie S4: WT-GCaMP 0.05% SDS

Movie S5: WT-GCaMP 5 mM H_2_O_2_

Movie S6: Empty-vector 5 mM H_2_O_2_

Movie S7: *cap1*Δ-GCaMP 5 mM H_2_O_2_

Movie S8: *hog1*Δ-GCaMP 5 mM H_2_O_2_

Movie S9: *cnb1*Δ-GCaMP 5 mM H_2_O_2_

Movie S10: *crz1*Δ-GCaMP 5 mM H_2_O_2_

Movie S11: *cch1*Δ-GCaMP 5 mM H_2_O_2_

Movie S12: *yvc1*Δ-GCaMP 5 mM H_2_O_2_

---

**Fig. S1:**
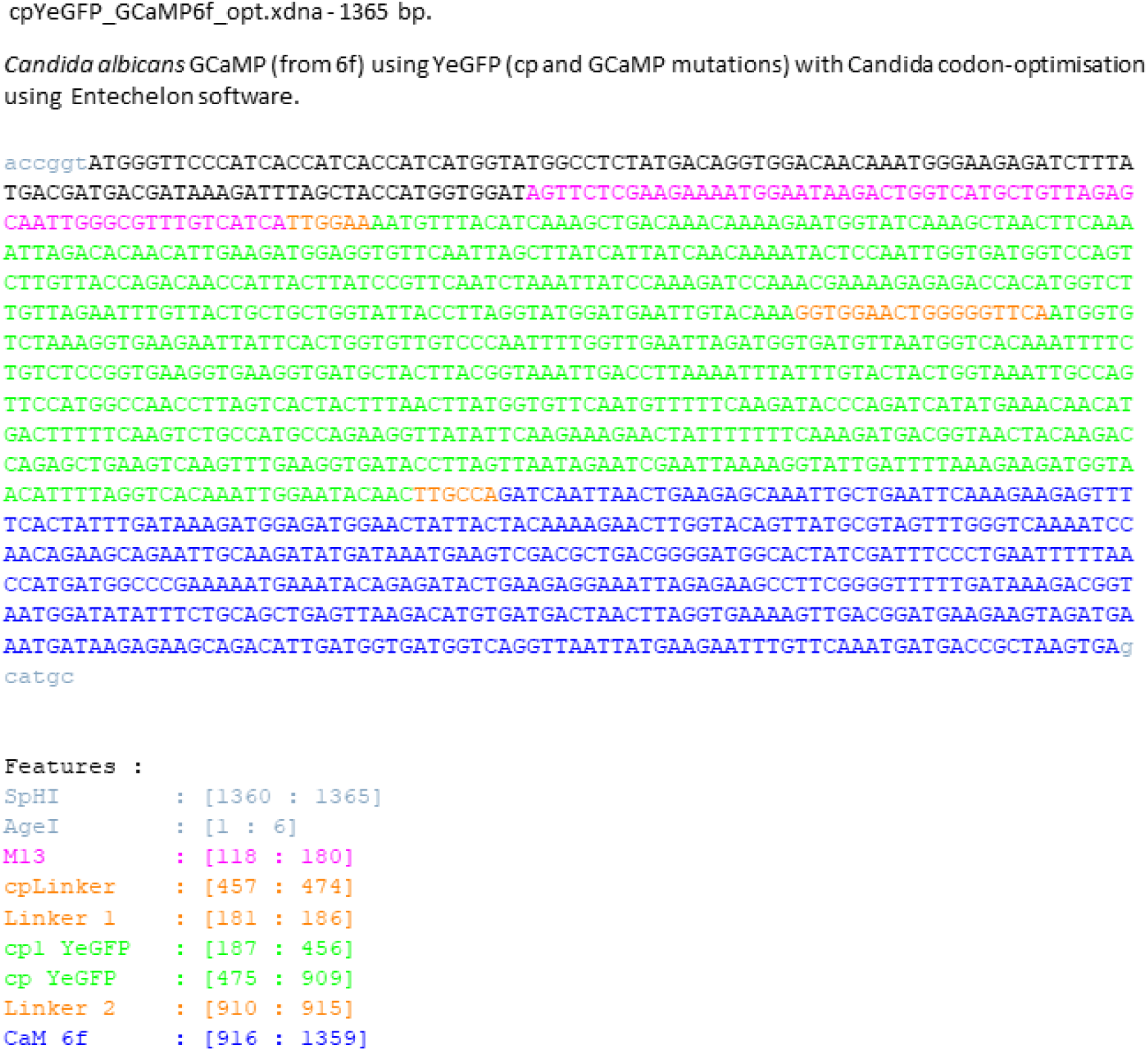
Codon optimised nucleotide sequence of GCaMP6f for expression in *C. albicans*. GCaMP6f coding sequence was taken from Chen, et al., 2013, codon optimised for expression in *C. albicans* and synthesised in plasmid Ca-GCaMP6f by Geneart. Features of coding sequence are highlighted in the figure.

**Fig. S2:**
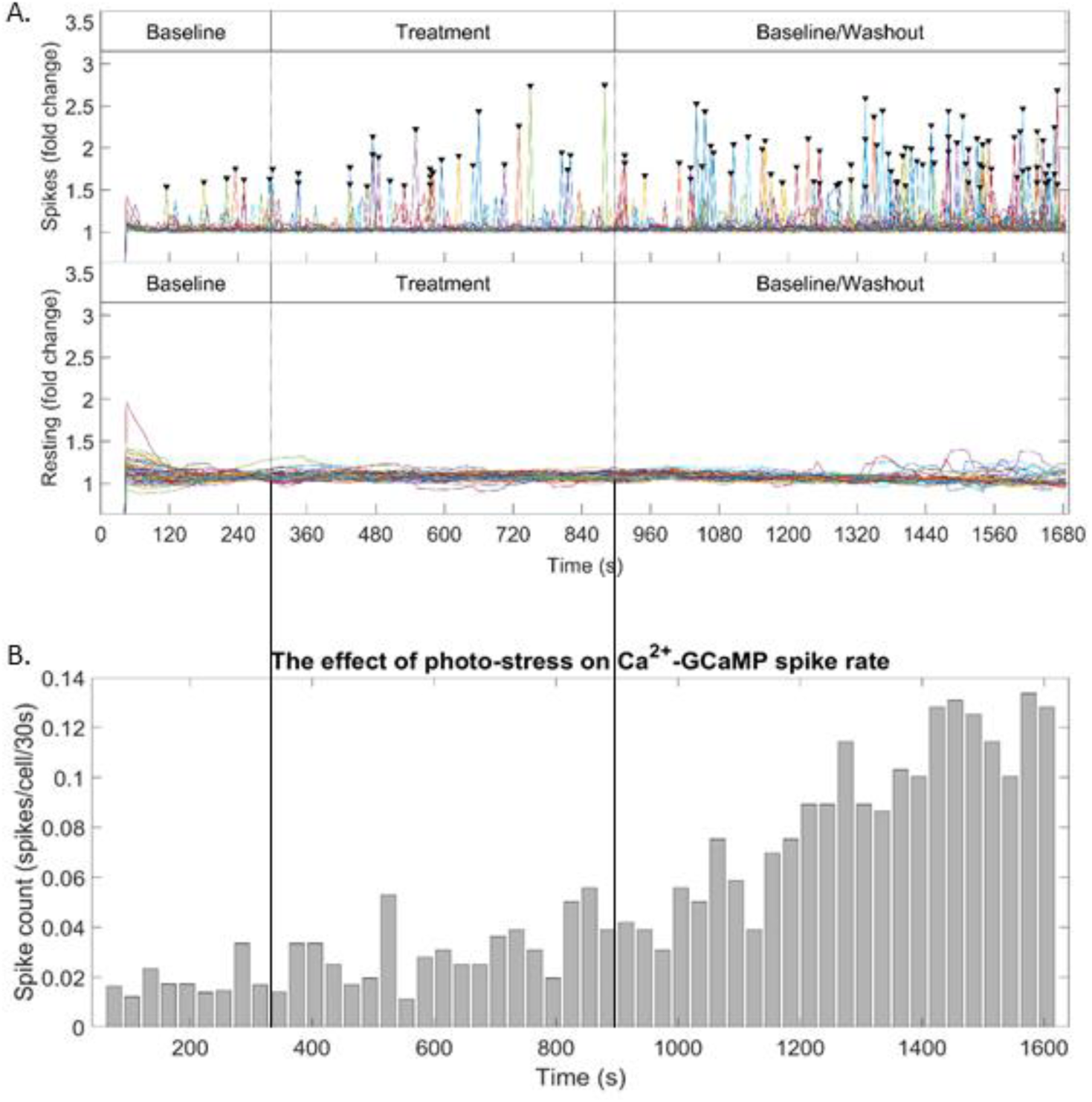
Onset of photo-stress occurs in Stage 3 (washout) of the experimental time-course. A. Representative GCaMP Ca^2+^ spiking and ambient Ca^2+^ GCaMP plots of BWP17-GCaMP in the presence of 5 mM Ca^2+^ (pH 7.5) throughout the entire experimental time course. B. Spikes/Cell/ 30 s window were plotted over the same time course.

**Fig. S3:**
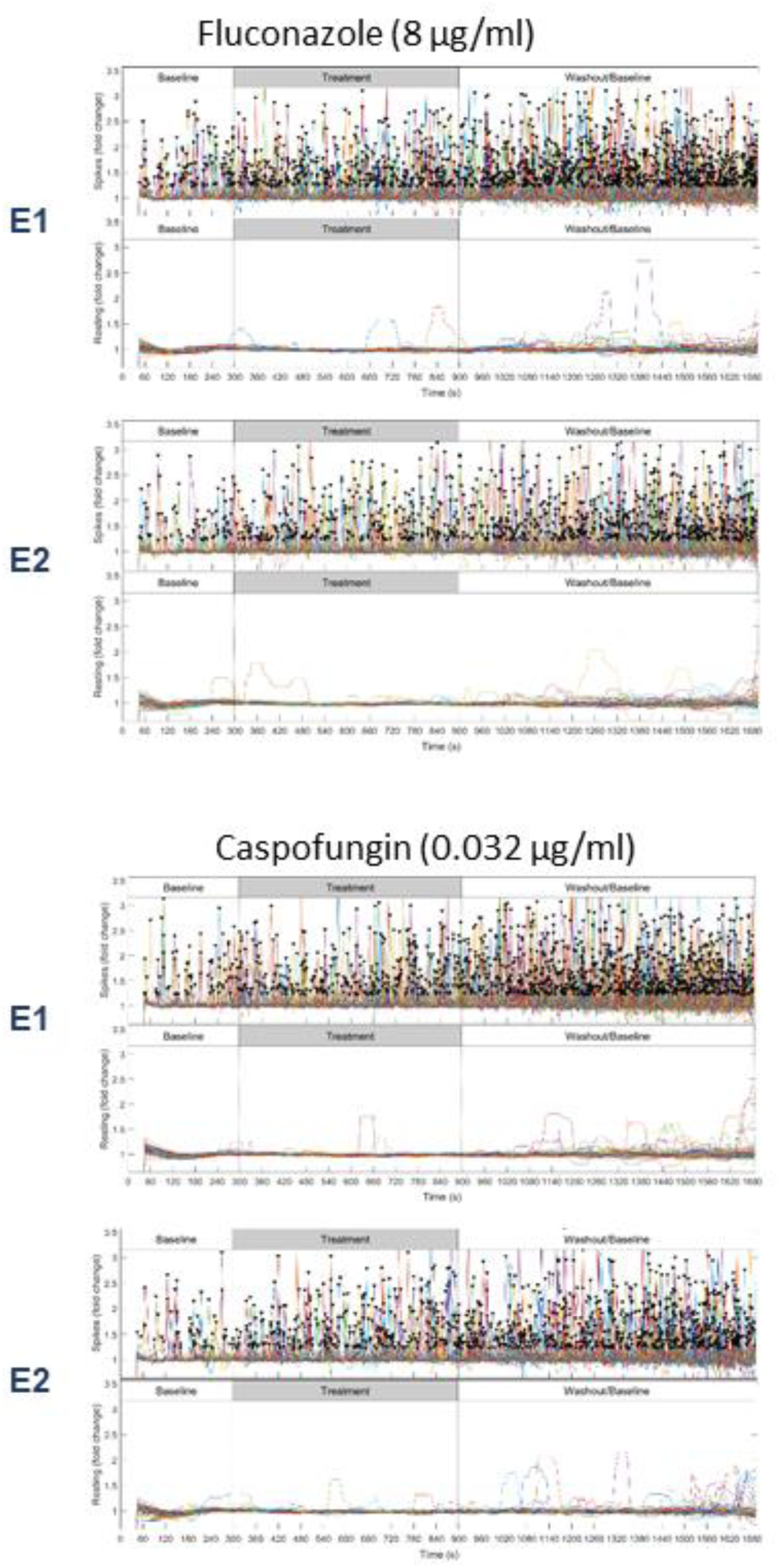
10 min exposures to fluconazole or caspofungin does not affect Ca^2+^ GCaMP activity. Ca^2+^ GCaMP and ambient Ca^2+^ GCaMP distributions of wild-type cells expressing GCaMP exposed twice to 8 ug/ml fluconazole or 0.032 ug/ml caspofungin during S2.

**Fig. S4:**
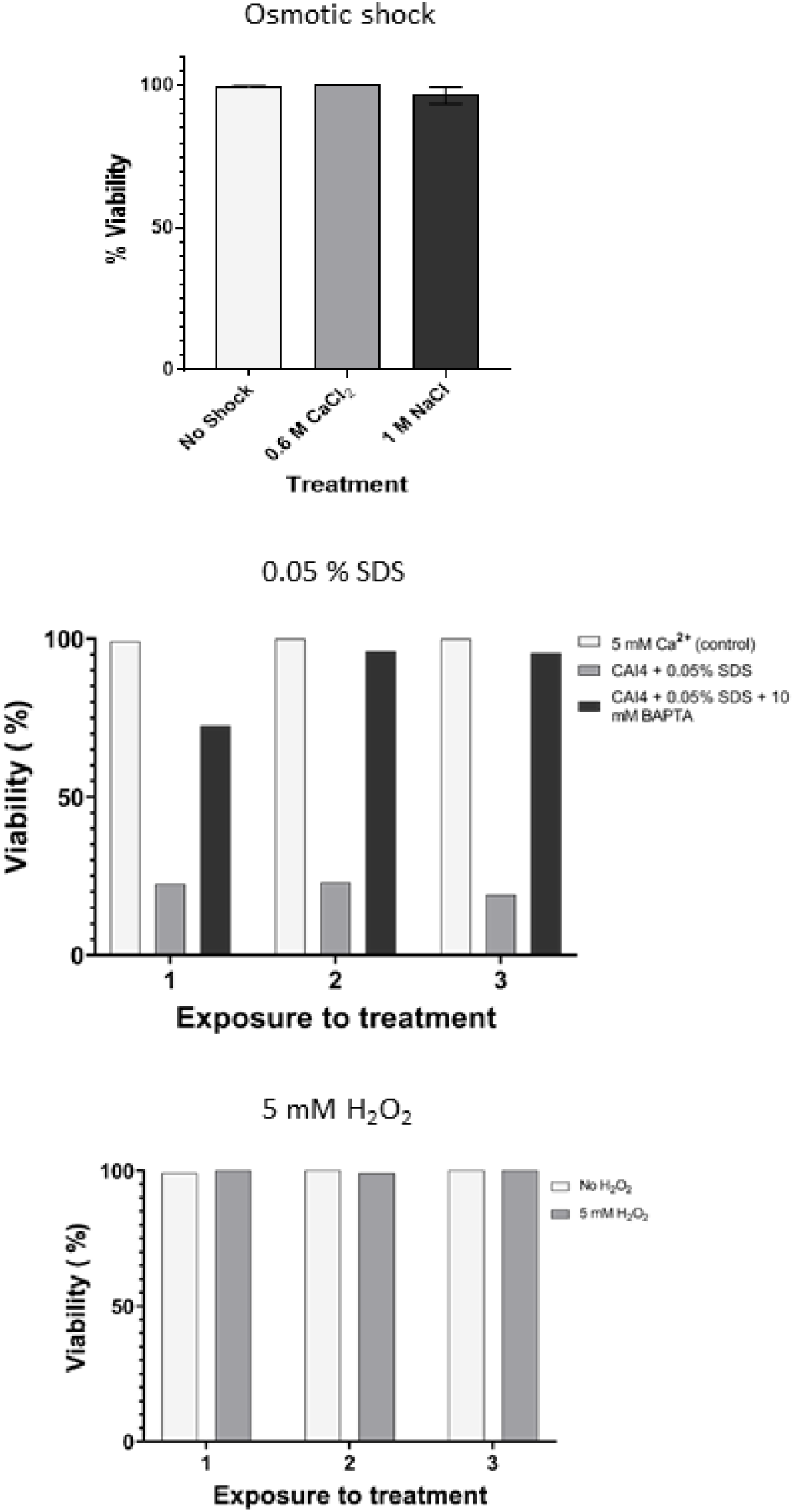
Cell viability of control strain expressing GCaMP after 3 exposures to osmotic shock, SDS, or H_2_O_2_. A. The empty vector strain (A569) was exposed to either 0.6 M CaCl_2_ or 1 M NaCl (5 mM Ca^2+^) 3 times during S2 and stained with 1 ug/ml PI at the end of E3. % viability was determined as the percentage of non stained cells in the population. The no shock control was BWP17-GCaMP in the presence of 5 mM Ca^2+^ and viability was determined as above. Bars = mean ± SD across 3 exposures B. CAI4-GCaMP was exposed 3 times to 0.05 % SDS (in 5 mM Ca^2+^) or 0.05 % SDS + 10 mM BAPTA (in 5 mM Ca^2+^) in S2 of the experimental time course, and % viability was calculated as in (A). C. BWP17-GCaMP was exposed 3 times to 5 mM H_2_O_2_ (in 5 mM Ca^2+^) during S2 of the experimental time course and % viability was calculated as in (A).

**Fig. S5:**
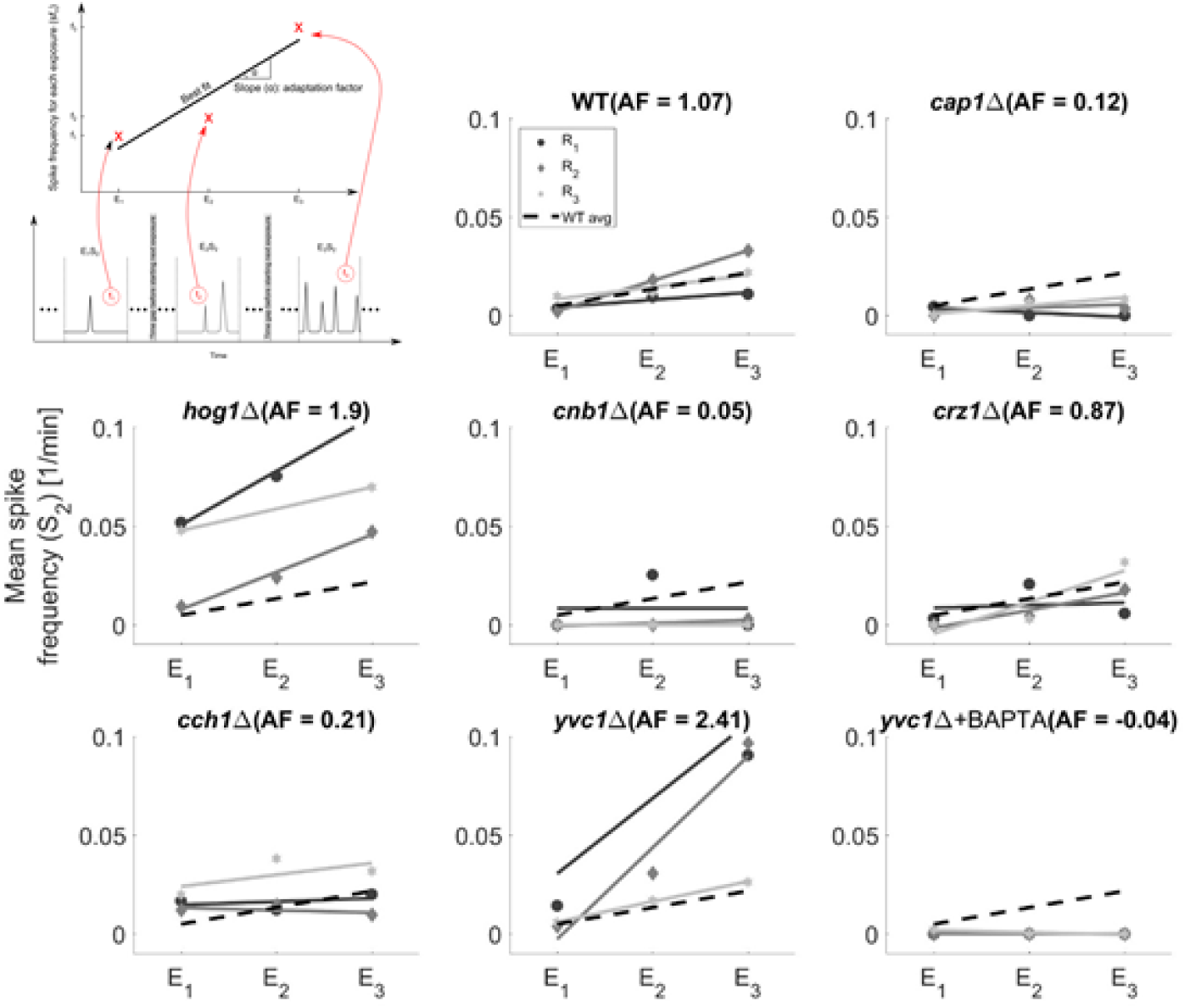
Adaptation Factors for WT and mutant strains treated with 5 mM H_2_O_2_. The adaptation factor was defined as slope of the change in spike frequency per cell from exposure 1 to exposure 3 (see illustration in figure). Plots showing the spike rate (spike/cell/minute) during S2 (exposure to H_2_O_2_) of WT and mutant strains. Lines represent line of best fit across each exposure. Each line represents individual biological replicate. Dotted line = mean WT value of 3 biological replicates. Values of mean spike frequency used in to determine the adaptation factor are shown in Table S1.

**Table S1:** Mean Ca^2+^-GCaMP spike frequency

| Strain Name | Rep1 | Rep2 | Rep3 | Mean + SD |
| --- | --- | --- | --- | --- |
| Wild-type | 1.07 | 1.53 | 0.61 | $1.07 \pm 0.46$ |
| *cap1Δ* | -0.22 | 0.18 | 0.41 | $0.12 \pm 0.32$ |
| *hog1Δ* | 2.72 | 1.89 | 1.10 | $1.90 \pm 0.81$ |
| *cnb1Δ* | 0 | 0.15 | 0 | $0.05 \pm 0.08$ |
| *crz1Δ* | 0.13 | 0.89 | 1.59 | $0.87 \pm 0.73$ |
| *cch1Δ* | 0.17 | -0.12 | 0.59 | $0.21 \pm 0.36$ |
| *yvc1Δ* | 3.81 | 2.41 | 1.02 | $2.41 \pm 1.39$ |
| *yvc1Δ*+BAPTA | 0 | 0 | -0.12 | $-0.04 \pm 0.07$ |

**Table S2:**
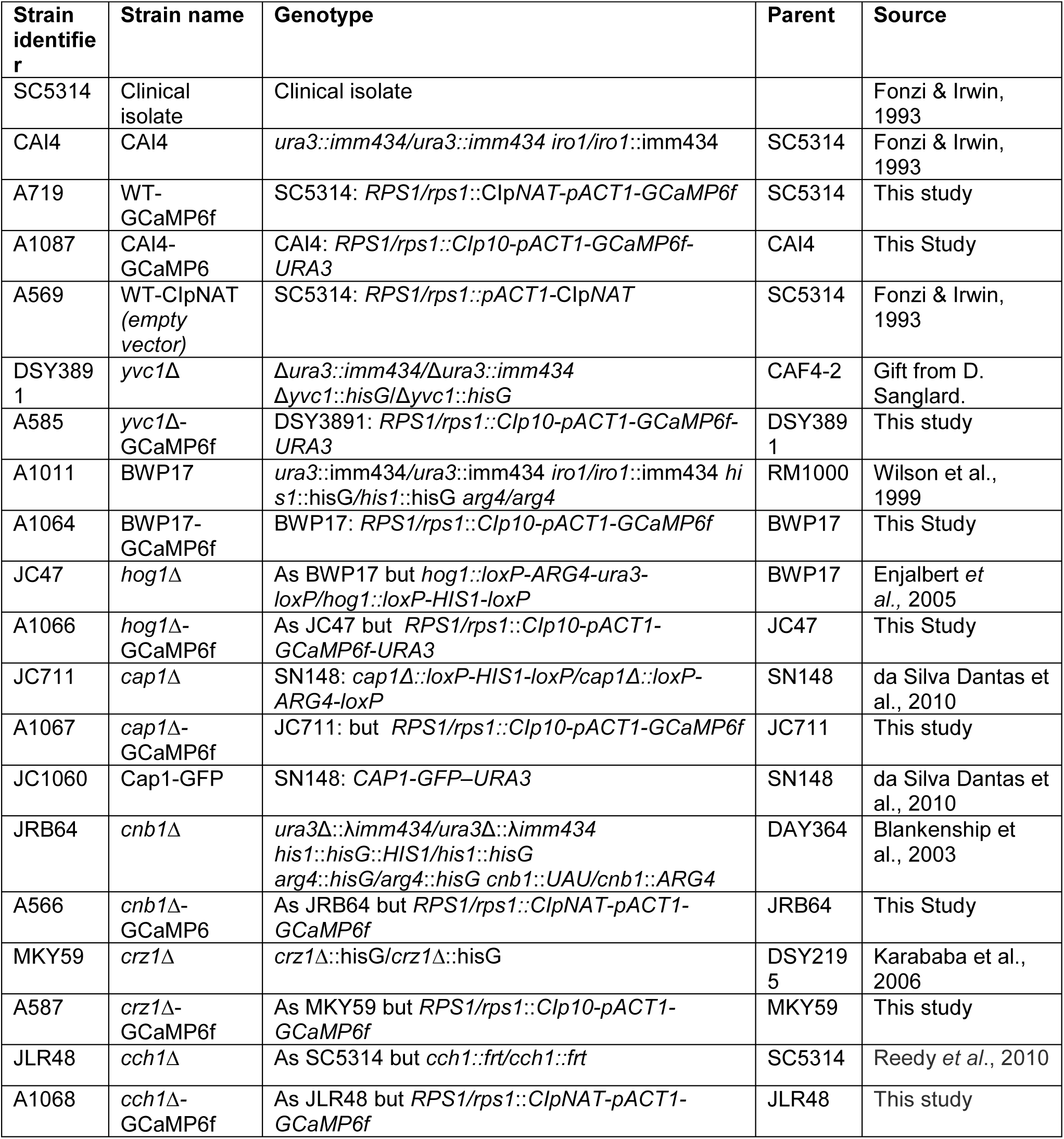
C. albicans strains used in this study

**Table S3:** Plasmids used in this study

| Plasmid name | Strain identifier | Description | Reference/Source |
| --- | --- | --- | --- |
| Clp10-*pACT1-ScCycT* |  | *Candida* integrating plasmid with selectable marker: *URA3* | Barelle *et al*, 2004 |
| Clp-NAT | B250 | Clp10 with NAT replacing *URA3* selection marker | Shahana *et al*, 2014 |
| Ca-GCaMP6f | B151 | Circularly-permuted, codon-optimized GCaMP6f | GeneArt<br>Chen *et al*, 2013 |
| ClpNAT-GCaMP6 | B153 | *ClpNATflip-ACT1p-GCaMP6f-ScCycT* | This study |
| Clp10-GCaMP6 | B251 | *Clp10-ACT1p-GCaMP6f-ScCycT (URA3)* | This study |

